# Different mechanisms of synapsin-induced vesicle clustering at inhibitory and excitatory synapses

**DOI:** 10.1101/2023.03.20.533583

**Authors:** Sang-Ho Song, George J. Augustine

## Abstract

Synapsins cluster synaptic vesicles (SVs) to provide a reserve pool (RP) of SVs that maintains synaptic transmission during sustained activity. However, it is unknown how synapsins cluster SVs. Here we show that either liquid-liquid phase separation (LLPS) or tetramerization-dependent cross-linking can cluster SVs, depending upon whether a synapse is excitatory or inhibitory. Cell-free reconstitution revealed that both mechanisms can cluster SVs, with tetramerization bring more effective. At inhibitory synapses, perturbing synapsin-dependent LLPS impairs SV clustering and synchronization of GABA release, while perturbing synapsin tetramerization does not. At glutamatergic synapses, the opposite is true: synapsin tetramerization enhances clustering of glutamatergic SVs and mobilization of these SVs from the RP, while synapsin LLPS does not. Comparison of inhibitory and excitatory transmission during prolonged synaptic activity revealed that synapsin LLPS serves as a brake to limit GABA release, while synapsin tetramerization enables rapid mobilization of SVs from the RP to sustain glutamate release.

## INTRODUCTION

One of the defining characteristics of chemical synapses is a dense cluster of synaptic vesicles (SVs) within their presynaptic terminals. How SVs are clustered remains unclear. One important clue has come from studies of synapsins, a family of SV-binding proteins that have been identified as key regulators of SV trafficking. Synapsins arise from 3 genes in mammals and have numerous presynaptic roles, including regulation of the kinetics of SV fusion with the plasma membrane (Hilfiker et al., 1998; Humeau et al., 2001; Hilfiker et al., 2005; Medrihan et al., 2013) and participation in various forms of short-term synaptic plasticity (Cheng et al., 2018). The best-established function of synapsins is to regulate the storage and mobilization of SVs within a reserve pool (RP), where they can be mobilized to replenish SVs consumed during synaptic activity (Landis et al., 1988; Hirokawa et al., 1989; Hilfiker et al., 1998; Hilfiker et al., 1999; Zhang and Augustine, 2021). However, the mechanism by which synapsins cluster SVs is unresolved.

Two main hypotheses have been proposed to account for the ability of synapsins to cluster SVs (Zhang and Augustine, 2021). One is based on the observation that SVs are connected by filamentous structures (Landis et al., 1988; Hirokawa et al., 1989; Siksou et al., 2007; Fernandez-Busnadiego et al., 2010; Cole et al., 2016; Wesseling et al., 2019) which are less numerous in mice lacking synapsins 1 and 2 (Wesseling et al., 2019). Further, given that synapsins bind to both SVs (Schiebler et al., 1986; Benfenati et al., 1989; Orlando et al., 2014) and to other synapsin molecules – as dimers (Hosaka and Sudhof, 1999; Orlando et al., 2014) or tetramers (Brautigam et al., 2004) - it is possible that such interactions can cross-link SVs to create the filamentous structures (Siksou et al., 2007; Orlando et al., 2014; Wesseling et al., 2019; Zhang and Augustine, 2021). However, there has been no definitive proof that synapsin oligomers are responsible for SV clustering (Zhang and Augustine, 2021). Alternatively, membrane-less condensates, formed by liquid-liquid phase separation (LLPS) (Shin and Brangwynne, 2017), can create subcellular synaptic structures (Zeng et al., 2016; Milovanovic et al., 2018; Wu et al., 2020). Synapsin 1a has been shown to form such condensates via LLPS; these structures accumulate small lipid vesicles *in vitro* (Milovanovic et al., 2018) and may be involved in SV clustering at synapses (Pechstein et al., 2020; Wu et al., 2020).

To discriminate between these two hypotheses, we used structure-function analyses, cell-free reconstitution, and physiological measurements of SV trafficking and transmitter release to revisit the LLPS and oligomerization properties of synapsins. Our conclusion is that synapsin LLPS plays an important role at inhibitory synapses, while synapsin oligomerization is important at excitatory synapses.

## RESULTS

### All synapsin isoforms generate liquid phase separation

In vertebrates, alternative splicing generates at least 5 major isoforms from the 3 synapsin genes: synapsins 1a/1b, 2a/2b, and 3a (Sudhof et al., 1989; Hosaka and Sudhof, 1998; Kao et al., 1998). All these isoforms contain an intrinsically disordered region (IDR) in their conserved C domains that is thought to be important for LLPS (Milovanovic et al., 2018). Because each isoform has a unique IDR (Figure S1), we asked whether differences in IDR length affect the ability of each synapsin isoform to produce LLPS. Purified synapsins 1a/1b, 2a/2b and 3a– tagged with enhanced green fluorescence protein (EGFP) – were found to form droplets *in vitro*, indicating LLPS (Figure 1A). Droplet size varied for each synapsin isoform (Figure 1B), with the isoform with the longest IDR isoform, synapsin 1a, forming the largest droplets, while synapsin 2b, with the shortest IDR, formed the smallest droplets (Figure 1C). For all isoforms, there was a strong correlation (*Pearson’s r* = 0.99) between IDR length and droplet diameter, indicating the importance of IDR length for producing LLPS (Figure 1D).

**Figure 1.**
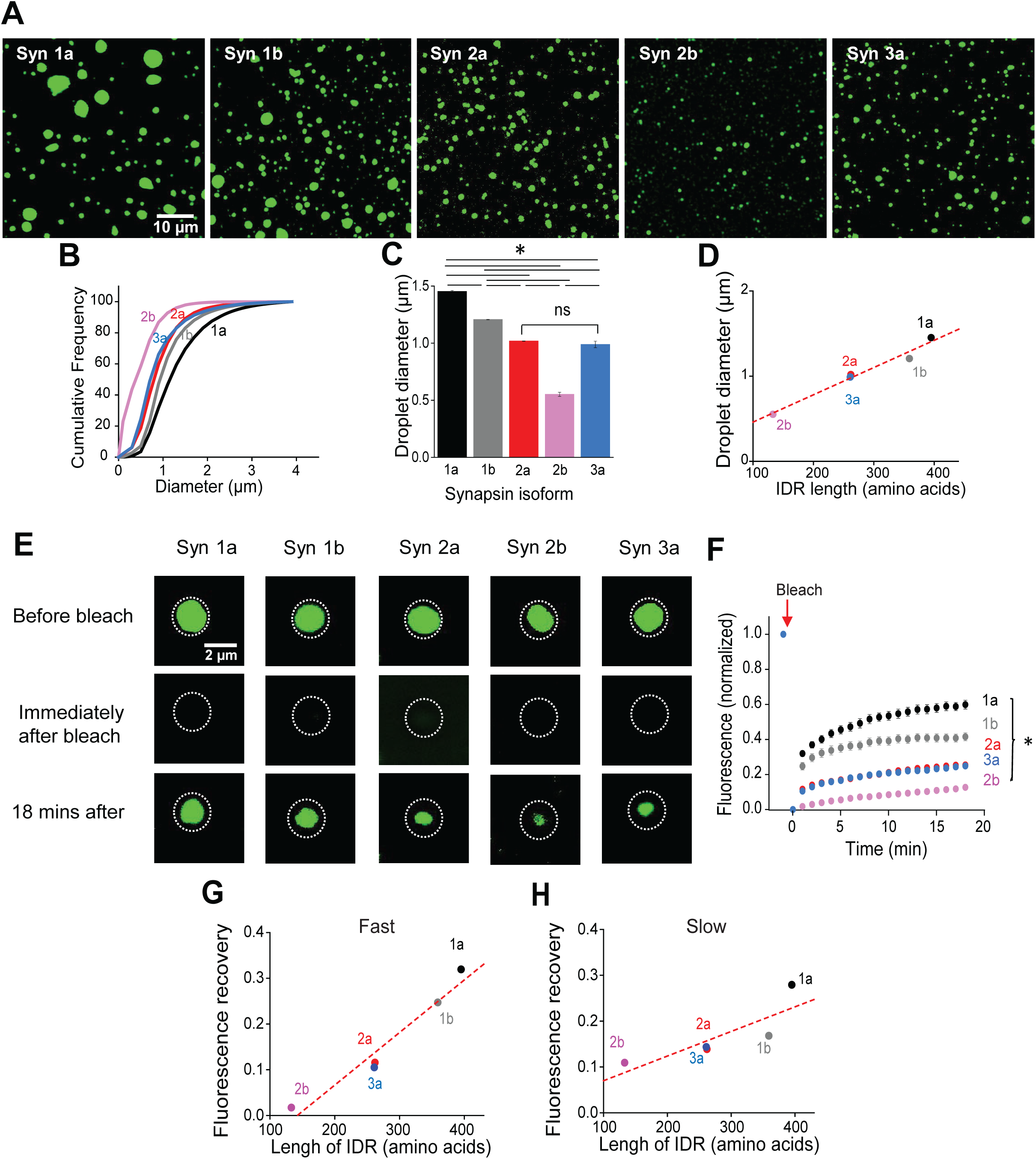
Liquid-liquid phase separation (LLPS) properties of synapsin isoforms. (A) Droplet formation by equivalent concentrations (10 µM) of EGFP-synapsin 1a, 1b, 2a, 2b and 3a. (B) Cumulative distribution of diameters of droplets formed by each of the 5 synapsin isoforms. (C) Mean diameters of synapsin droplets. Number of independent replicates ranged from 3 to 5. Every comparison between isoforms show significant differences except synapsin 2a and 3a. (D) Linear correlation between droplet diameter and IDR length of synapsin isoforms. (E) Images of synapsin droplets before and after photobleaching. (F) Time course of fluorescence recovery after bleaching for each synapsin isoform. Number of replicates ranged from 17-28. (G) Linear correlation between rate of initial, fast component of fluorescence recovery (measured at 1 min after bleaching) and synapsin IDR length. (H) Linear correlation between rate of slow component of fluorescence recovery, measured between 1 and 20 min after bleaching, and synapsin IDR length. Statistical comparisons in (C) and (F) were done with ANOVA, followed by Tukey’s post-hoc test; asterisks indicate significant differences (p<0.05).

To measure the ability of synapsin isoforms to diffuse to/from LLPS-induced droplets, we used fluorescence recovery after photobleaching (FRAP). For each synapsin isoform, droplets of similar size (∼ 2 μm diameter) were completely bleached and we measured the subsequent recovery of fluorescence, caused by diffusion of EGFP-tagged synapsin molecules from outside the droplet (Figures 1E and 1F). Fluorescence recovery was observed for all synapsin isoforms and consisted of two components: a rapid initial phase, complete within 1 minute, that likely represents incorporation of freely-diffusing unbleached synapsin molecules, as well as a slower phase that was largely complete within 20 minutes. The latter reflects incorporation of synapsin from nearby unbleached droplets, either by gradual replacement or by fusion of smaller unbleached droplets. For both components, the degree of recovery was highly correlated (*Pearson’s r* = 0.95) with isoform IDR length (Figures 1G and 1H). Thus, the LLPS properties of synapsin isoforms are determined by their IDR length.

### Two mechanisms of SV clustering by synapsin 2a *in vitro*

LLPS formation by synapsins and other proteins is affected by ATP, a cellular energy source that is also a biological hydrotrope (Milovanovic et al., 2018). We examined the effects of ATP on LLPS produced by synapsin 2a, the only isoform that rescues the physiological phenotype at glutamatergic synapses of synapsin triple knock-out (TKO) mice (Gitler et al., 2008). ATP impaired LLPS droplet formation (Figure 2A). Half-maximal inhibition of synapsin 2a droplet fluorescence occurred at 1.7 mM ATP (Figure 2B), within the physiological range of intracellular ATP concentration (1-2 mM) (Rangaraju et al., 2014).

**Figure 2.**
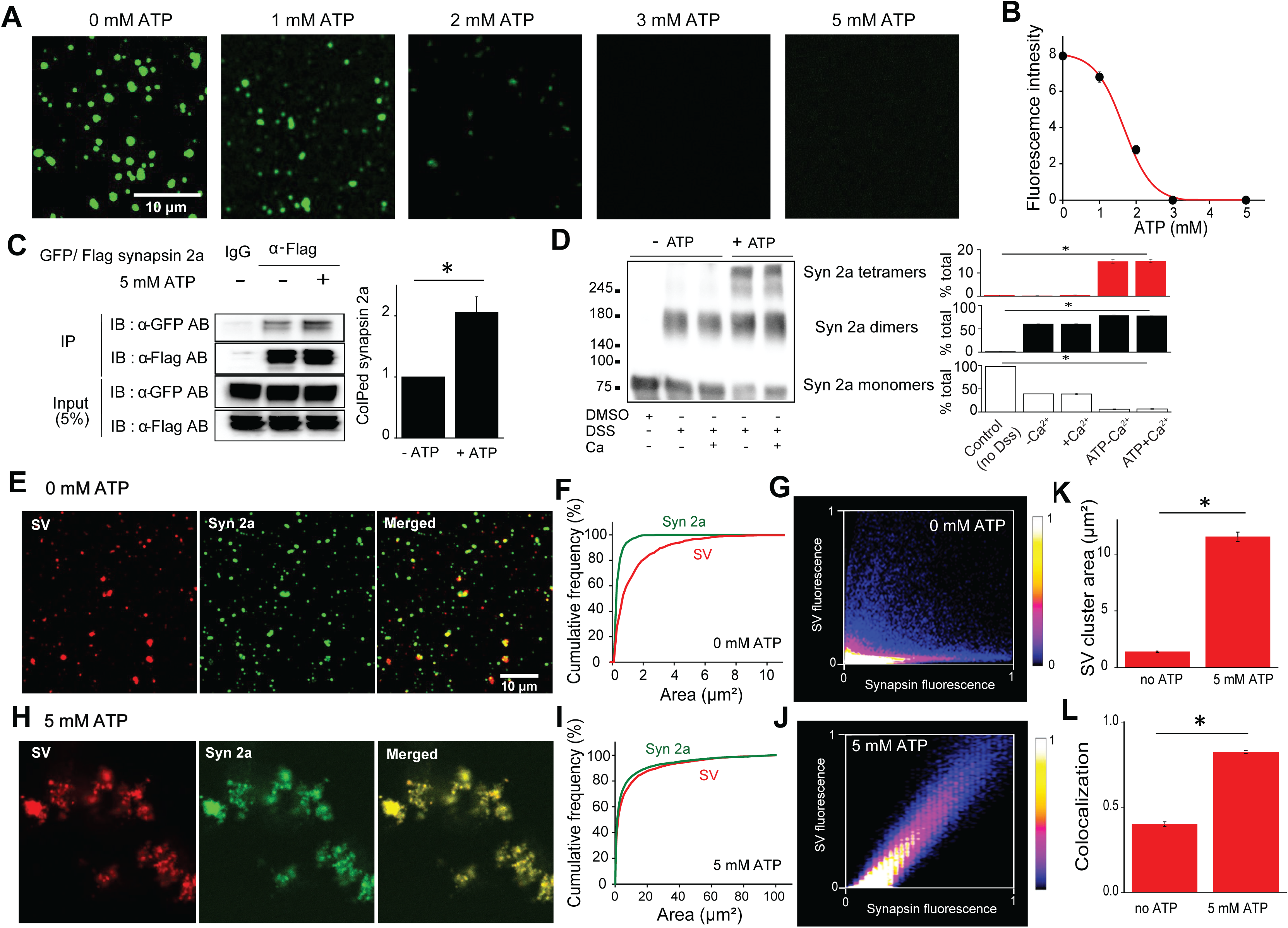
Synaptic vesicle clustering via synapsin 2a LLPS and oligomerization. (A) Fluorescence images of droplets formed by EGFP-synapsin 2a LLPS at different ATP concentrations. Images are representative of 30 fields from 3 replicates. (B) Relationship between droplet fluorescence and ATP concentration. Values represent mean ratio of fluorescence intensity inside/outside of droplets. (C) Effect of ATP on synapsin oligomerization, measured via immunoprecipitation. Two tagged forms of synapsin 2a (GFP, Flag) were expressed in HEK 293T cells and cell lysates were incubated with the indicated antibodies. ATP (5 mM) increased the amount of pull-down of synapsin 2a, indicating increased oligomerization. (D) Identification of synapsin 2a oligomeric forms. Purified synapsin 2a was incubated in the presence of ATP and/or Ca^2+^ and was subjected to crosslinking by DSS. *Left*, Western blot of synapsin monomers, dimers and tetramers in indicated conditions. *Right*, ATP (5 mM) increased the amount of synapsin dimers and tetramers, while Ca^2+^ (2.1 mM) had no effect. Statistical comparisons in (C) and (D) were done with ANOVA, followed by Tukey’s post-hoc test; asterisks indicate significant differences (p<0.05; n=4). (E-L) *In vitro* reconstitution of synaptic vesicle clustering, under conditions favoring synapsin LLPS (0 mM ATP) or oligomerization (5 mM ATP). (E) When purified GFP-synapsin 2a and DiI-stained SVs were combined in the absence of ATP, both SV clusters (red) and synapsin 2a (green) formed punctate structures. Merged image (right) compares location of SVs and synapsin 2a; overlapping pixels are indicated in yellow. (F) Cumulative distribution of areas of synapsin 2a droplets and SV clusters; note difference in sizes of these two structures. (G) Scatter plot of SV fluorescence and synapsin fluorescence in individual pixels in images similar to those shown in (E). (H) The presence of ATP (5 mM) caused synapsin 2a and SVs to form much larger structures; merged images (right) indicated the high degree of co-localization of these structures. (I) Similar sizes of synapsin 2a structures and SV clusters; note 10-fold difference in x-axis scale compared to (F). (J) High degree of correlation between SV fluorescence and synapsin fluorescence in individual pixels in images similar to those shown in (H). (K) Mean areas of SV clusters produced by synapsin 2a in the absence and presence of ATP. (L) Comparison of colocalization, measured by the Pearson’s correlation coefficient, between SVs and synapsin 2a SV in the absence and presence of ATP. Data in (E) to (L) are representative of 47-54 images from 3 replicates. Statistical comparisons in (K) and (L) were done with a t-test; asterisks indicate significant differences (p<0.05).

ATP also increases the dimerization (Hosaka and Sudhof, 1999) and tetramerization (Brautigam et al., 2004) of synapsin 1a. We found that 5 mM ATP enhanced cross-binding of two tagged versions of recombinant synapsin 2a (Figure 2C). To determine whether ATP produced synapsin 2a dimers or tetramers, a chemical crosslinker (DSS) was employed(Orlando et al., 2014). Purified, his-tagged synapsin 2a was incubated in the presence or absence of ATP, as well as with or without Ca^2+^ (2.1 mM), which also affects oligomerization (Hosaka and Sudhof, 1999; Orlando et al., 2014). In the absence of ATP, synapsin 2a largely dimerized; ATP enhanced both dimer and tetramer formation, with nearly no synapsin 2a in an unbound state at 5 mM ATP (Figure 2D). Further, Ca^2+^ did not affect this equilibrium. Thus, the absence of ATP favors LLPS by synapsin 2a, while ATP promotes oligomerization.

These observations indicate that ATP can identify the relative contributions of LLPS and oligomerization to the ability of synapsin 2a to cluster SVs *in vitro*. To reconstitute SV clustering, synapsin-free SVs were purified from TKO mouse brains (Figure S2B) and mixed with purified EGFP-synapsin 2a. In the absence of ATP, synapsin 2a had some ability to cluster SVs (Figure 2E). SVs clusters were larger than synapsin 2a droplets (Figure 2F), suggesting that not all SV clusters were associated with droplets. Indeed, the fluorescence of labelled SVs only loosely colocalized with that of synapsin 2a, indicated by yellow pixels in the merged image of Figure 2E. Pixel-by-pixel comparisons indicated a mild correlation between SV and synapsin 2a signals (Figure 2G). In contrast, when 5 mM ATP was added, SV clusters were approximately 10-times larger (Figures 2H and 2I; note 10-fold change in x-axis scale from Figure 2F) and very similar in size to clusters of synapsin 2a fluorescence (Figure 2I). SV and synapsin 2a fluorescence signals were tightly colocalized, indicated both by yellow pixels in merged images (Figure 2H) and in the relationship between the fluorescence of these two signals within each pixel (Figure 2J). Thus, when LLPS was eliminated and oligomerization was maximized by 5 mM ATP, SV clusters were larger (Figure 2K) and colocalization between SVs and synapsin 2a was greater (Figure 2L). Our interpretation is that oligomerization of synapsins bound to adjacent SVs cross-links the SVs, thereby yielding large SV clusters and a higher degree of colocalization. In summary, both LLPS and synapsin oligomerization can cluster SVs, with oligomerization quantitatively more effective *in vitro*.

### Synapsin 2a mutations that impair oligomerization but not liquid phase separation

While ATP is a valuable tool for evaluating LLPS and oligomerization *in vitro*, manipulating ATP levels in live cells would disrupt many other cellular processes. We therefore developed an alternative approach for cellular analyses by using synapsin 2a mutations that affect ATP binding or oligomerization. We identified a point mutant, synapsin 2a K337Q, that disrupts a lysine important for hydrogen bonding involved in tetramerization (Esser et al., 1998; Brautigam et al., 2004), as well as a second mutant, synapsin 2a K270Q, that reduces ATP binding (Shulman et al., 2015). Both mutant proteins exhibited less self-association, both in the absence or presence of ATP (Figure S2A). Chemical cross-linking showed that while both mutants formed dimers, neither synapsin 2a mutant formed tetramers in the presence of ATP (Figure 3A). In contrast, LLPS was little affected by either mutation (Figure 3B): K270Q generated slightly larger droplets than synapsin 2a WT, while K337Q formed slightly smaller droplets (Figures 3B and 3C). Similar results were observed in FRAP measurements of protein mobility (Figures S3A-S3C).

**Figure 3.**
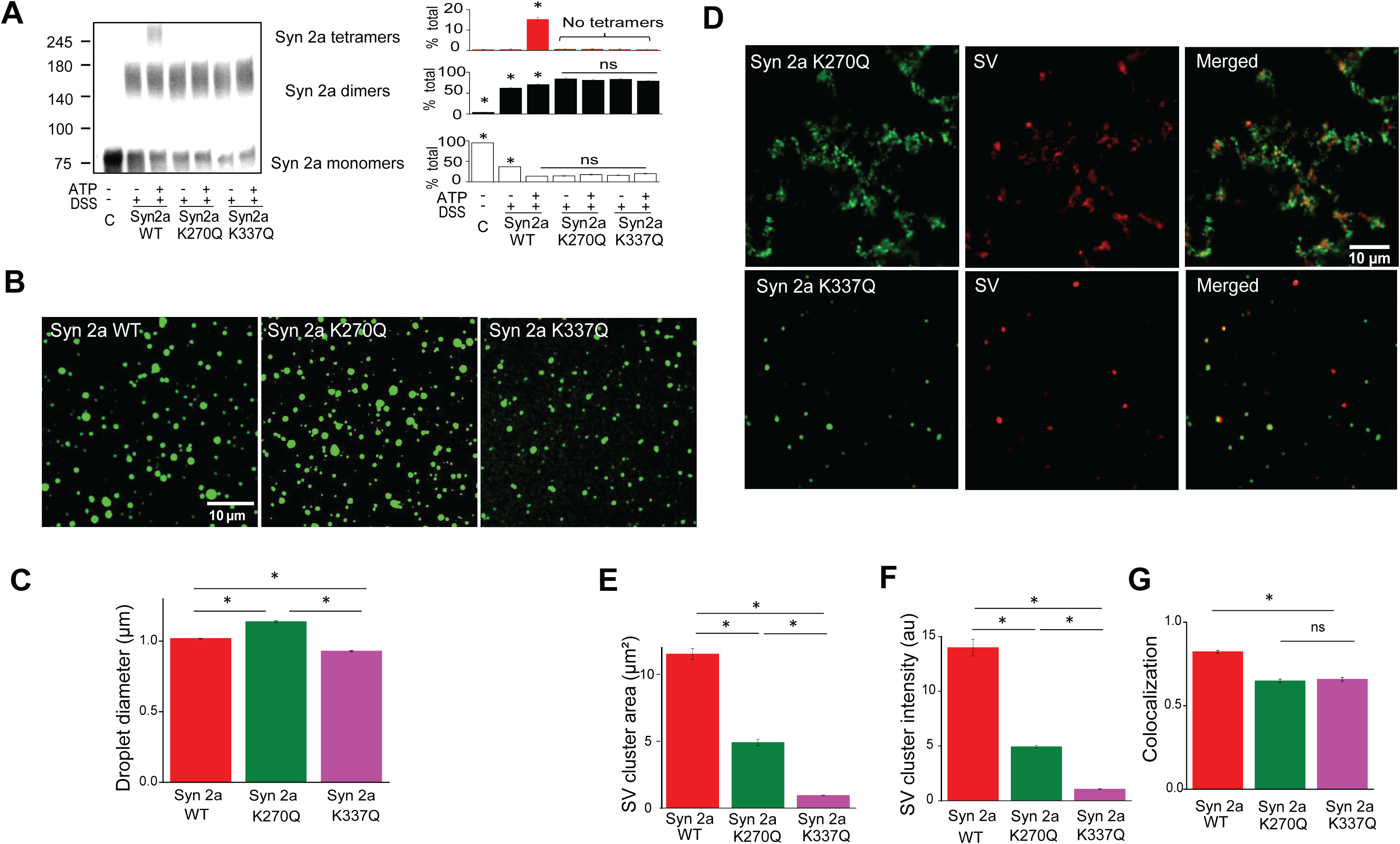
Synapsin 2a mutants that disrupt tetramerization and SV clustering. (A) Impairment of synapsin 2a tetramerization *in vitro* by K270Q and K337Q mutations. *Left*, Western blot of synapsin monomers, dimers and tetramers in indicated conditions. *Right*, while ATP (5 mM) increased the amount of synapsin dimers and tetramers for wild-type synapsin 2a (WT), only dimers and monomers were observed for K270Q and K337Q mutant proteins. (B) LLPS-induced droplets for EGFP-synapsin 2a WT, K270Q and K337Q. (C) Mean diameters of synapsin droplets in images such as those in (C). Statistical comparisons in (A) and (C) were done with ANOVA, followed by Tukey’s post-hoc test; asterisks indicate significant differences (p<0.05; n=4). (D) SV clusters (red) and synapsin structures (green) produced by EGFP-synapsin 2a K270Q and K337Q in the presence of 5 mM ATP. Both K270Q and K337Q reduced the mean area of SV clusters (E) mean fluorescence intensity of SV clusters (F) and degree of SV/synapsin colocalization, measured by the Pearson’s correlation coefficient (G). Number of images ranged from 45-63 for each condition, from 3 independent replicates.

Compared to WT synapsin 2a (Figure 2H), both mutants had a reduced capacity to cluster SVs in the presence of 5 mM ATP (Figure 3D). In particular, SV clustering was barely apparent with the K337Q mutant protein (Figure 3D, lower). The area of SV clusters was significantly reduced with K270Q and nearly abolished with K337Q (Figure 3E and 3F). Further, the degree of colocalization of SVs and synapsin 2a was reduced for both mutants in comparison to WT (Figure 3G). These differences are not due to changes in the ability of the synapsin 2a mutants to bind to SVs (Figures S3D-S3H). These results provide additional support for our conclusion that the large SV clusters observed in the presence of ATP rely on synapsin oligomerization – specifically tetramerization - and resultant SV cross-linking.

### Liquid phase separation supports SV clustering and the reserve pool of inhibitory synapses

We next evaluated the importance of synapsin tetramerization and LLPS in trafficking of SVs at synapses by reexpressing synapsin 2a or expressing synapsin 2a mutants in cultured hippocampal neurons from synapsin-free (TKO) mice. Because previous work has shown that synapsins have different functions at inhibitory and excitatory synapses (Gitler et al., 2004a; Song and Augustine, 2016), we considered each separately.

For both types of synapses, the fluorescent dye, FM 4-64, was used to determine the size of SV pools (Chi et al., 2001; Mozhayeva et al., 2002; Gitler et al., 2008), followed by posthoc immunocytochemical labelling to identify glutamatergic (VGlut, Figure S5A) and GABAergic (VGAT) synapses (Figure 4A). For inhibitory synapses, the readily-releasable pool (RRP) was defined from the amount of FM4-64 released by a brief stimulus (10 Hz, 10 s, Figure S4A), while the dye released by subsequent exhaustive stimulation (10 Hz, 470 s) reflected reserve pool (RP) size (Figure 4B). The total SV pool was defined as the sum of the RRP and RP. At inhibitory synapses, the two synapsin 2a mutants defective in tetramerization increased RP size in comparison to the RP of TKO neurons not expressing any synapsins, as did wild-type synapsin 2a (Figure 4C). In contrast, RRP size was unaffected by synapsin expression (Figure 4D). As a result, the total SV pool was larger in TKO neurons expressing any synapsin 2a variant (Figure 4E). This result indicates that the RP of GABAergic SVs is maintained in the absence of synapsin tetramerization.

**Figure 4.**
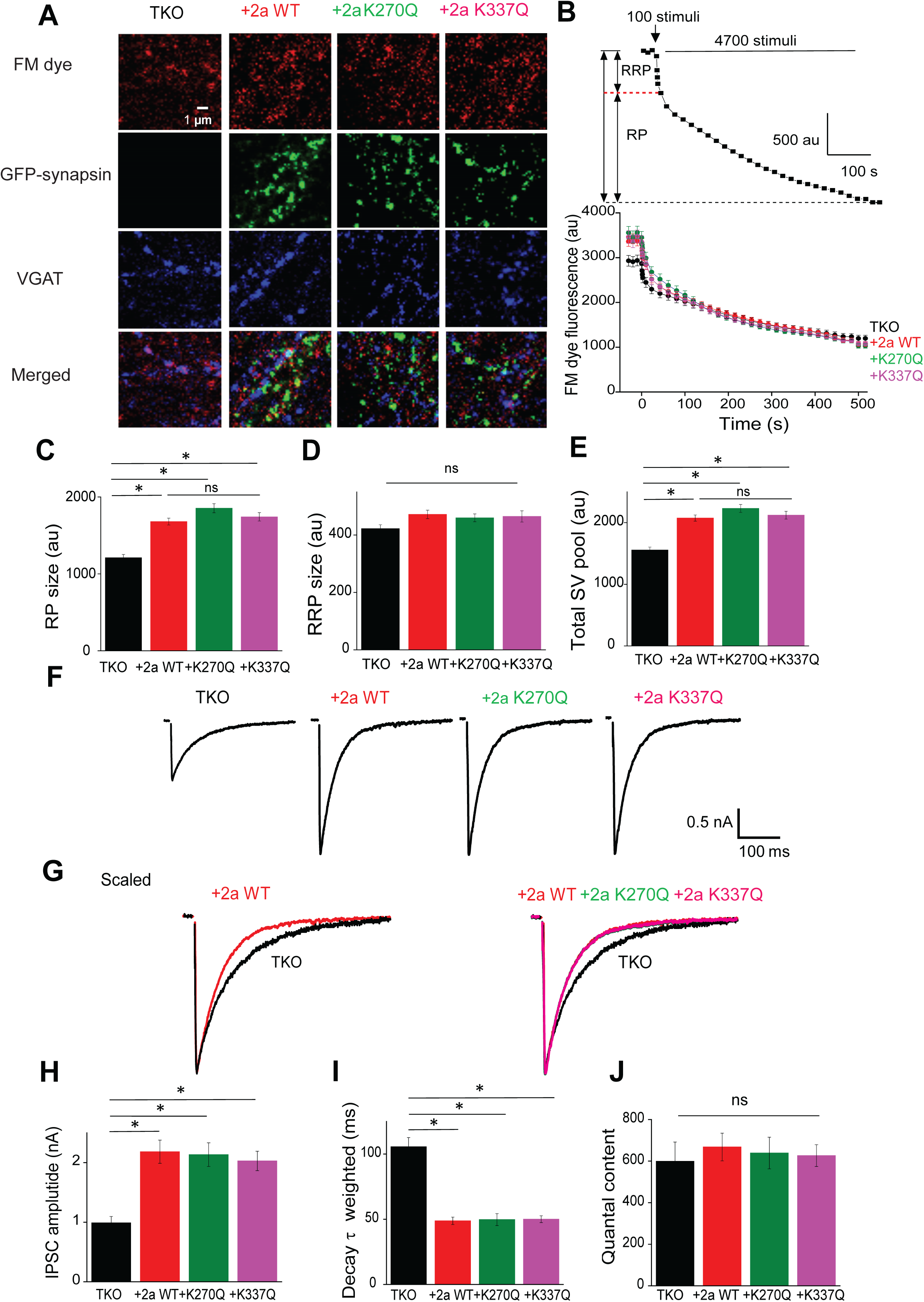
Synapsin 2a tetramerization does not regulate GABA release at inhibitory synapses. (A) Representative images of FM4-64 dye loading (top row) of presynaptic terminals expressing GFP-synapsin 2a (second row). Post-hoc immunostaining for VGAT (third row) indicates GABAergic presynaptic terminals. Bottom row is merger of images in 3 rows above, to compare co-localization of fluorescent signals. (B) Kinetics of FM dye destaining. *Top*, representative plot of destaining time course illustrates depletion of the readily releasable pool (RRP) by an initial 100 stimuli, followed by a prolonged loss of dye from the reserve pool (RP) during sustained stimulation (all at 10 Hz). *Bottom*, comparison of magnitude and time course of FM4-64 destaining in TKO neurons expressing either no synapsins or one of the 3 synapsin 2a variants indicated. (C-E) Effects of synapsin 2a variants on SV pools, measured from FM4-64 destaining results such as those shown in (B). All synapsin 2a variants increased RP size (C) and total SV pool size (E), but had no effect on RRP size (D). Statistical comparisons were done with ANOVA, followed by Tukey’s post-hoc test; asterisks indicate significant differences (p<0.05). Number of synapses analyzed ranged from 98 to 134. (F-J) Measurements of GABA release evoked by presynaptic action potentials, in TKO neurons expressing either no synapsins or one of the synapsin 2a variants. (F) Representative inhibitory postsynaptic currents (IPSCs) from TKO neurons in the 4 indicated conditions. (G) Superimposed IPSC were scaled to the same peaks to illustrate the kinetics of IPSC decay. All synapsin 2a variants, including synapsin 2a WT (*left*) and the two tetramerization-defective mutants (*right),* rescued the slowing of IPSC decay kinetics observed in TKO neurons. As a result of the synchronization of GABA release shown in (G) synapsin 2a WT and both tetramerization- deficient synapsin 2a mutants rescued the mean peak amplitude of IPSCs (H), the time constant of IPSC decay (I), and IPSC quantal content (J). Statistical comparisons in (H) to (J) were done with ANOVA, followed by Tukey’s post-hoc test; asterisks indicate significant differences (p<0.05). Number of cells used to generate data ranged from 14-17.

To identify the contributions of synapsin LLPS and tetramerization to synaptic transmission, we measured inhibitory synaptic responses in cultured autaptic neurons from the hippocampus of TKO mice. At such synapses, loss of synapsins desynchronizes GABA release, leading to a reduction in the peak amplitude of inhibitory postsynaptic currents (IPSCs) evoked by presynaptic action potentials and a slowing of IPSC decay (Gitler et al., 2004a; Song and Augustine, 2016). Confirming a previous report (Song and Augustine, 2016), expression of synapsin 2a WT in TKO neurons rescued this effect, both increasing IPSC amplitude (Figure 4F) and accelerating IPSC kinetics (Figure 4G, left). Similar effects were found in TKO neurons expressing either synapsin 2a mutant (Figures 4F and 4G). There were no significant differences in the ability of these synapsin 2a variants to rescue IPSC amplitude (Figure 4H) or rate of decay (Figure 4I). Similarly, the quantal content – a measure of how many SVs are discharged in response to a presynaptic action potential - was constant in all conditions (Figure 4J). There was also no difference in spontaneous GABA release between all groups, as measured by miniature IPSCs (Figures S4B-S4E). In summary, GABA release evoked by presynaptic action potentials (Del Castillo and Katz, 1954; Heuser et al., 1979) was rescued by all synapsin 2a variants, regardless of their ability to undergo tetramerization. These results indicate that synapsin tetramerization, and resultant cross-linking of GABA-containing SVs, is unnecessary for GABA release under physiological conditions. Because the synapsin 2a variants do undergo LLPS, this suggests that LLPS may be required for trafficking of GABA-containing SVs.

### SV cross-linking is important at glutamatergic synapses

We next examined the function of synapsin LLPS and oligomerization at excitatory synapses, where synapsins maintain a RP that is mobilized during repetitive activity: at TKO glutamatergic synapses, basal synaptic transmission is normal, but depresses more rapidly during repetitive activity (Gitler et al., 2004a; Gitler et al., 2008). Although all synapsin isoforms have extensive IDRs and undergo LLPS (Figure 1), synapsin 2a is the only isoform that rescues the synaptic depression phenotype of glutamatergic synapses (Gitler et al., 2008). As a result, there is no correlation between the IDR length of these isoforms and their previously reported ability to rescue synaptic depression (Figure 5A). Notably, synapsin 1a - which has the longest IDR (Figure S1F) and produces the most robust LLPS (Figure 1D) – does not rescue the glutamatergic synaptic depression phenotype (Gitler et al., 2008). This suggests that a mechanism other than LLPS maintains the glutamatergic SV reserve pool.

**Figure 5.**
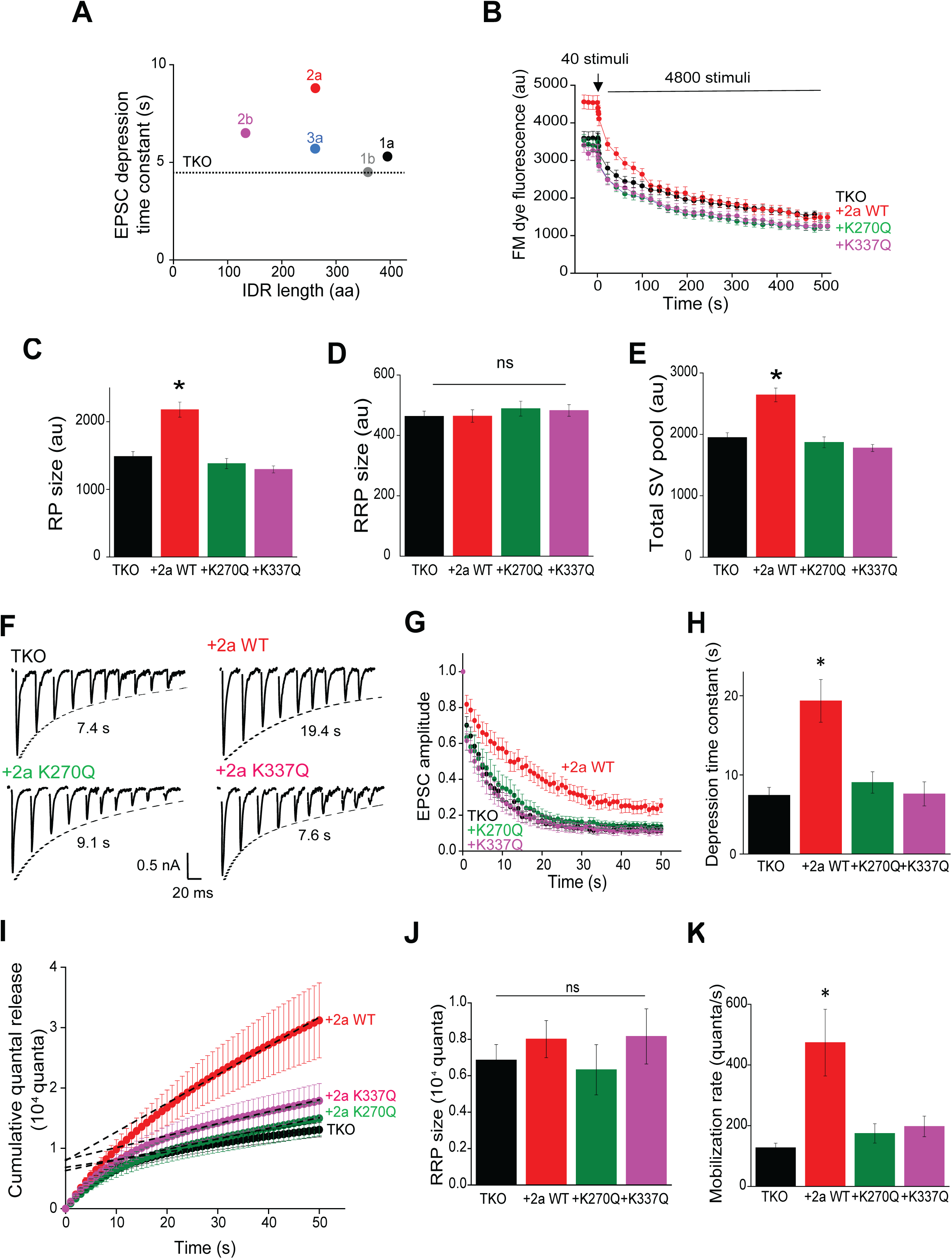
Role of synapsin 2a tetramerization in glutamate release at excitatory synapses. (A) Relationship between the ability of synapsin isoforms to rescue synaptic depression at glutamatergic neurons from synapsin TKO mice (Gitler et al., 2008) and isoform IDR length. Dashed line indicates the rate of depression observed in TKO neurons not expressing any synapsins (i.e. no rescue). (B) FM4-64 dye destaining in TKO neurons expressing either no synapsins (black) or the indicated synapsin 2a variants. Dye destaining induced by the first 40 stimuli unloaded SVs in the RRP, while the subsequent 4800 stimuli (all at 10 Hz) unloaded dye from the RP. (C to E) Effects of synapsin 2a variants on SV pools, measured from FM4-64 destaining results such as those shown in (B). Only synapsin 2a WT could rescue RP size (C) and total SV pool size (E); synapsins had no effect on RRP size (D). Statistical comparisons in (C) to (E) were done with ANOVA, followed by Tukey’s post-hoc test; asterisks indicate significant differences (p<0.05). Number of excitatory synapses used to generate data in (B) to (E) ranged from 88 to 136. (F) Representative excitatory postsynaptic currents (EPSCs; every 50^th^ response) during trains of stimuli (10 Hz) recorded in TKO neurons expressing either no synapsins (TKO) or the indicated synapsin 2a variants. (G) Time course of depression of EPSC amplitude during stimuli similar to those shown in (F). EPSC amplitude was normalized to first response in the train. (H) Mean time constant of depression, determined from exponential fits to data shown in (B). Only synapsin 2a WT rescued the synaptic depression phenotype of TKO neurons. (I) Time course of cumulative EPSC charge, obtained by integrating responses shown in (G). Data from 20-50 s were fitted by linear regression; dotted lines indicate extrapolation back to y-intercept, to estimate RRP size. Slope of these lines indicates the rate of mobilization (J) Mean RRP size was synapsin-independent. (K) Mean SV mobilization rates for indicated conditions; only synapsin 2a WT rescued the deficit in SV mobilization observed in TKO neurons. Statistical comparisons in (H), (J) and (K) were done with ANOVA, followed by Tukey’s post-hoc test; asterisks indicate significant differences (p<0.05). Number of cells used to generate data in (G) to (K) ranged from 14 to 17.

FM 4-64 measurements of pool sizes at glutamatergic synapses (Figure 5B and Figure S5A) indicated that expression of synapsin 2a WT increased the size of the RP above that observed in TKO neurons, while neither K270Q nor K337Q increased RP size (Figure 5C). None of the synapsin variants affected RRP size (Figure 5D). Thus, only synapsin 2a WT rescued the RP phenotype of excitatory TKO neurons; this indicates that both the total number of glutamatergic SVs, as well as the number of SVs in the RP, depends on synapsin oligomerization. To consider the role of synapsin tetramerization in excitatory synaptic transmission, we expressed synapsin 2a WT, K270Q or K337Q in cultured TKO neurons. Basal synaptic transmission, measured as the quantal content of excitatory postsynaptic currents (EPSCs) evoked by single action potentials, was similar in all conditions (Figures S5B and S5C). However, the rate of synaptic depression evoked by trains of action potentials (10 Hz, 50 s) varied for the different synapsin 2a variants (Figure 5F). Depression was accelerated in TKO neurons but was rescued by expression of synapsin 2a WT (Figure 5G) (Gitler et al., 2008) . In contrast, TKO neurons expressing either tetramerization-deficient form of synapsin 2a exhibited rapid depression (Figures 5F and 5G). Quantitative comparison indicated that the time constant of depression was rescued significantly only by synapsin 2a WT (Figure 5H).

We also determined whether synapsin 2a oligomerization influences the size of the RRP or the rate of mobilization of glutamatergic vesicles from the RP by analyzing the total amount of glutamate released during trains of action potentials (Schneggenburger et al., 1999; Stevens and Williams, 2007). The amount of transmitter released during a train of stimuli exhibited two components: a rapid initial phase, associated with depletion of the RRP, followed by a slower, linear increase that reflected SV mobilization from the RP (Figure 5I). Extrapolating the slow component back to its y-intercept defined RRP size (dashed lines in Figure 5I), which was unchanged in all conditions (Figure 5J). The rate of mobilization of SVs from the RP to the RRP was determined by the slope of the late component of integrated eEPSC charge: while synapsin 2a significantly enhanced the rate of mobilization (Gitler et al., 2008), neither tetramerization-defective mutant rescued the mobilization rate (Figure 5K).

Our FM 4-64 and electrophysiological analyses of excitatory synapses are consistent in indicating that synapsin tetramerization and resultant SV cross-linking is important both for maintaining the RP of glutamatergic terminals and for mobilization of SVs from this pool. The fact that the synapsin 2a mutants, as well as other synapsin isoforms, support LLPS (Figures 1C and 3C) but not glutamatergic RP size (Figure 5C) or mobilization from the RP (Figure 5K) suggests that LLPS does not play a role in these processes.

### Perturbing synapsin LLPS selectively impairs GABAergic synapses

The results above indicate different roles for synapsin LLPS at inhibitory and excitatory synapses. To further test this idea, we examined the actions of the SH3A domain of intersectin, which inhibits the ability of synapsins to undergo LLPS (Pechstein et al., 2020) by binding to a proline-rich region within the IDR (Gerth et al., 2017; Pechstein et al., 2020). While the SH3A domain inhibited synapsin LLPS *in vitro* (Figures 6A and 6B), it did not affect synapsin 2a tetramerization (Figure S6). As a negative control, we observed that the SH3B domain of intersectin, which does not bind to synapsins, influenced neither LLPS nor tetramerization (Figure 6A and Figure S6). At inhibitory synapses, the SH3A domain prevented synapsin 2a from rescuing the TKO phenotype: GABA release was desynchronized, evident as both a reduced peak amplitude of IPSCs evoked by presynaptic action potentials and a slowing of IPSC decay (Figures 6C-6E). In contrast, the SH3B domain did not interfere with the ability of synapsin 2a to rescue the TKO phenotype (Figures 6D-6F). These results indicate that disruption of synapsin LLPS impairs GABA release even though tetramerization is intact. At glutamatergic synapses, neither the SH3A domain nor the SH3B domain impaired the ability of synapsin 2a to rescue the depression phenotype of excitatory synapsin TKO neurons. This was evident as a slower rate of depression (Figures 6G and 6H) and higher mobilization rate (Figures 6I and 6J) compared to TKO. Thus, impairment of synapsin LLPS does not affect glutamate release but does impact GABA release.

**Figure 6.**
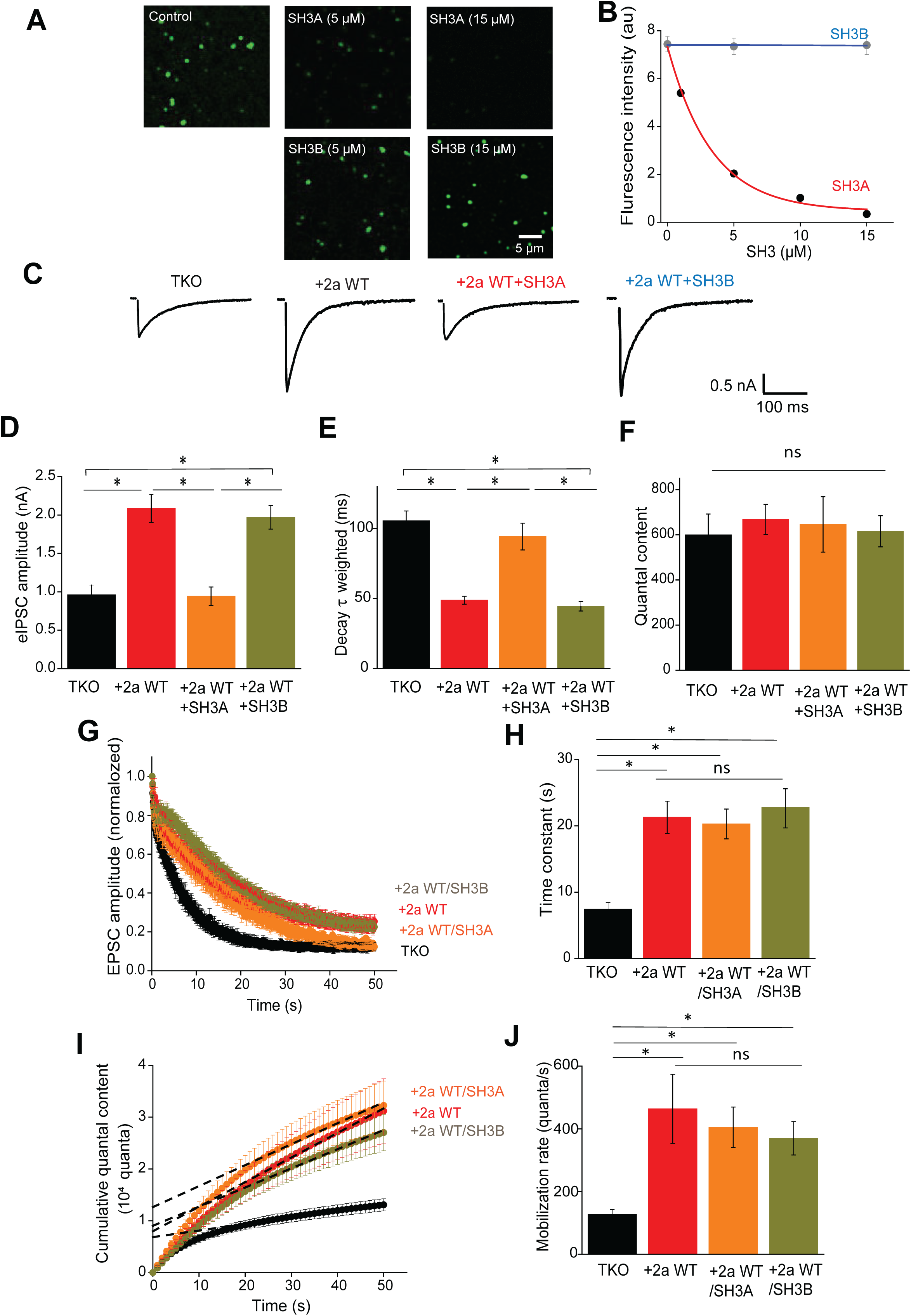
Perturbation of synapsin LLPS by intersectin SH3A domain selectively alters inhibitory synaptic transmission. (A) Fluorescence images of LLPS-induced droplets of EGF-synapsin 2a (1 µM) in the presence of mCherry-tagged intersectin SH3A (upper) or intersectin SH3B (lower). (B) Relationship between SH3 domain concentration and synapsin 2a-induced LLPS. Values represent mean ratio of fluorescence intensity inside/outside of synapsin 2a droplets. Number of images used ranged from 20 to 29, in 3 independent replicates. (C-F) IPSCs were selectively affected by intersectin SH3A domain. (C) Representative IPSCs from TKO neurons expressing no synapsins or synapsin 2a WT, with or without intersectin SH3A or SH3B domains. d, Mean amplitude of IPSC amplitudes measured in the indicated conditions; intersectin SH3A domain selectively blocked rescue of IPSC amplitude by synapsin 2a WT. (E) IPSC decay time constants (weighted double exponential fit) in the indicated conditions. Intersectin SH3A domain selectively blocked rescue of IPSC kinetics by synapsin 2a WT. (F) IPSC quantal content, determined by IPSC charge, was independent of synapsins. Statistical comparisons in (D) to (F) were done with ANOVA, followed by Tukey’s post-hoc test; asterisks indicate significant differences (p<0.05). Number of cells used in these measurements ranged from 14 to 17. (G-J) Depression of EPSCs at glutamatergic synapses is unaffected by intersectin SH3 domains. (G) Time course of depression of EPSC amplitude during stimulus trains (10 Hz); EPSC amplitude was normalized to first response in the train. (H) Mean time constant of depression, determined from exponential fits to data shown in (G). The ability of synapsin 2a WT to rescue the synaptic depression phenotype of TKO neurons was unaffected by intersectin SH3A or SH3B domains. (I) Time course of cumulative EPSC charge, obtained by integrating responses shown in (G). Data from 20-50 s were fitted by linear regression; dotted lines indicate extrapolation back to y-intercept, to estimate RRP size, while slope of these lines indicates the rate of SV mobilization. (J) Mean SV mobilization rates for indicated conditions; neither intersectin SH3A nor SH3B affected the ability of synapsin 2a WT to rescue the deficit in SV mobilization observed in TKO neurons. Statistical comparisons in (H) to (J) were done with ANOVA, followed by Tukey’s post-hoc test; asterisks indicate significant differences (p<0.05). Number of cells used to generate data ranged from 14 to 19.

### Synapsin tetramerization enables efficient mobilization of glutamatergic vesicles

To understand the logic behind employing two different synapsin-dependent mechanisms for SV clustering, we compared the mobilization of RP vesicles at GABAergic and glutamatergic synapses. RRP size and RP mobilization rate were determined from measurements of the cumulative number of quanta released during train stimulation (50 Hz for 50 s; Figure 7A). RRP size was similar between the two synapses (Figure 7B), consistent with a previous demonstration that the number of GABAergic and glutamatergic synapses is similar in microisland cultured hippocampal neurons (Chang et al., 2014). This is also consistent with our FM4-64 dye imaging, which showed that the RRPs of individual GABAergic and glutamatergic boutons are each approximately 400 fluorescence units when measured under identical conditions (Figures 4D, 5D and S7B). Remarkably, glutamatergic synapses exhibited a three-fold higher rate of SV mobilization from the RP to the RRP (Figure 7C). This higher rate of SV mobilization indicates that tetramerization-dependent SV crosslinking at glutamatergic synapses enables 3 times more SVs to be mobilized, per unit time.

**Figure 7.**
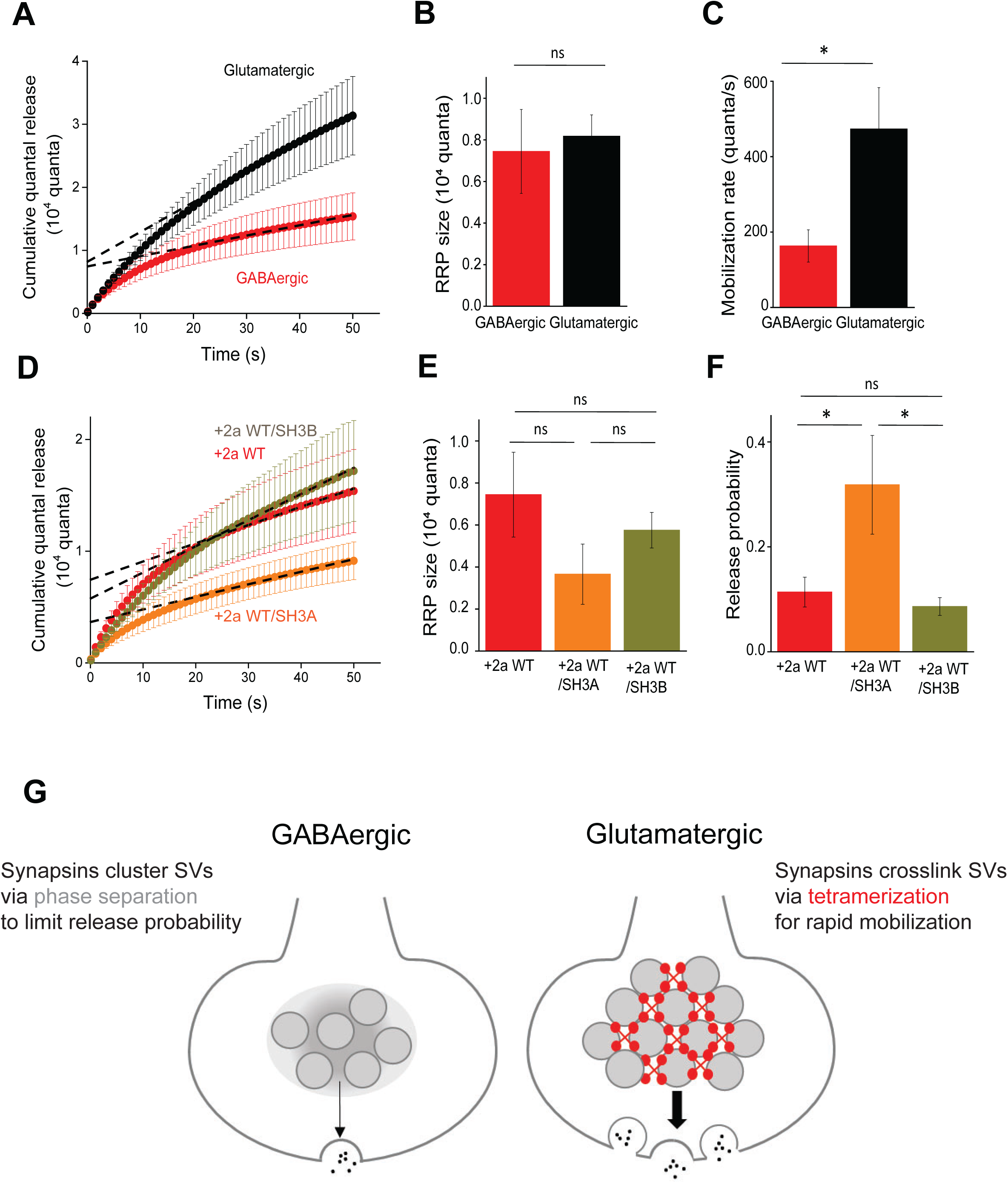
Comparison of transmission at glutamatergic and GABAergic synapses. (A) Time course of cumulative IPSC and EPSC charge from TKO neurons expressing 2a WT, obtained by integrating responses during train stimulation (50 Hz, 50 s) as shown in Figure 5I. Data from 20-50 s were fit by linear regression; dotted lines indicate extrapolation back to y-intercept, to estimate RRP size. Slope of these lines indicates the rate of mobilization. (B) Mean RRP size was comparable between the two different synapses. (C) Mobilization rates of glutamatergic synapses were larger than those of GABAergic synapses. Statistical comparisons in (B) and (C) were done with a t-test; asterisks indicate significant differences (p<0.05). Number of cells used to generate data was 10 (GABAergic) and 15 (glutamatergic). (D) Effects of perturbing synapsin-dependent LLPS on inhibitory transmission during train stimulation (50 Hz, 50 s) as in (A). (E) RRP size was not significant altered by interfering with LLPS via intersectin SH3A, or by the SH3B control. (F) GABA release probabilities were determined by the quantal content of individual IPSCs, divided by the number of quanta within the RRP; SH3A, but not SH3B, significantly increased release probability. Statistical comparisons in (E) and (F) were done with ANOVA with Tukey’s post hoc test; asterisks indicate significant differences (p<0.05), while *ns* indicates no significant difference. Number of cells used to generate data was 10 (+2a WT), 8 (+2a WT/SH3A) and 6 (+2a WT/SH3B). (G) Model of SV clustering and trafficking by synapsin LLPS and tetramerization. At GABAergic synapses, SVs are clustered by synapsin-dependent LLPS, while at glutamatergic synapses, synapsins crosslink SVs via tetramerization. Thickness of arrows indicates rate of mobilization of SVs from the RP, while the number of exocytotic events indicates transmitter release probability.

Faster SV mobilization also implies that the RP of SVs may be larger at glutamatergic synapses than at GABAergic synapses. This possibility was examined by comparing FM4-64 measurements of RP size in TKO neurons expressing synapsin 2a: mean RP size was indeed larger at glutamatergic boutons (2100 fluorescence units; Figure 5C and S7A), than at GABAergic boutons (1700 units; Figure 4C and S7A). To directly compare RP size and RP mobilization rates, we estimated the number of quanta in the RP. This estimation took advantage of our observation that RRP size of both GABAergic and glutamatergic synapses was consistent between FM4-64 and electrophysiological measurements. This allowed us to convert FM4-64 fluorescence units into SV quanta: RRP size measured by FM4-64 (Figures 4D, 5D and S7B) was divided by the number of quanta in the RRP, as determined electrophysiologically (Figure 7B). This defined the amount of FM4-64 destaining that corresponded to a quantum of glutamate or GABA, which was then used to convert FM4-64 measurements of RP size (Figure S7A) into number of quanta. With this transformation, we found that the number of quanta in the RP of glutamatergic synapses was larger than that of GABAergic synapses (Figure S7C). In summary, these analyses show that both the size of the RP (140%) and the rate of mobilization of SVs from the RP to the RRP (300%) are larger in glutamatergic synapses in comparison to GABAergic synapses. Thus, synapsin tetramerization enables greater mobilization of SVs from the RP during repetitive synaptic activity.

Next, to further define the role of synapsin LLPS in GABA release, we examined the effect of the SH3A domain on RP mobilization. RRP size and RP mobilization rate were determined by the slope of linear fit from the cumulative number of quanta released during train stimulation (50 Hz for 50 s; Figure 7D). Neither SH3A, nor the SH3B control, had a significant effect on RRP size (Figure 7E) or on the rate of SV replenishment from the RP to RRP (Figure S7D). Surprisingly, GABA release probability was significant increased when synapsin LLPS was perturbed by the SH3A domain (Figure 7F). All these results indicate that synapsin LLPS lowers the rate of SV mobilization from a smaller RP at GABAergic synapses, in comparison to glutamatergic synapses, and also reduces the probability of GABA release from the RRP. Thus, the role of synapsin LLPS is to maintain GABA release during repetitive synaptic activity, despite a smaller RP, by slowing consumption of GABA-filled SVs in the RRP.

## DISCUSSION

We have characterized LLPS and oligomerization of synapsins, have established that both of these processes can cluster SVs *in vitro*, and have identified distinct roles for LLPS and oligomerization in SV trafficking mediated by synapsin 2a. Our results indicate that synapsin LLPS, rather than cross-linking of GABA-containing SVs via synapsin tetramerization, is required for the RP of GABAergic SVs. In addition, retards LLPS release probability, which allows GABA release to be sustained from the smaller RP during repetitive activity. This contrasts with trafficking of glutamatergic SVs, where synapsin tetramerization provides a larger RP and faster mobilization of SVs within this pool, while LLPS does not play a detectable role (Figure 7G). Thus, although both LLPS and tetramerization contribute to SV trafficking, each mechanism makes unique contributions at hippocampal excitatory and inhibitory synapses, reflecting the different properties and functional requirements of these two types of synapses.

### LLPS and clustering of GABAergic SVs

We found that all five major synapsin isoforms can generate LLPS, with the size of droplets produced by LLPS and the mobile fraction measured by FRAP highly correlated with the IDR length of each isoform. These isoform-specific differences may have physiological consequences: multiple synapsin isoforms are expressed in individual neurons, with different isoform combinations in each type of presynaptic terminal (Wilhelm et al., 2014; Song and Augustine, 2015). It has been proposed that LLPS by synapsin can create a RP by separating the SV cluster from other presynaptic compartments (Milovanovic et al., 2018; Pechstein et al., 2020; Wu et al., 2020; Zhang and Augustine, 2021). We have found evidence supporting this when reconstituting SV clustering *in vitro* (Figure 2), measuring SV pool sizes optically (Figure 4) and monitoring inhibitory synaptic transmission (Figures 4, 6 and 7). Our conclusion that synapsin LLPS generates SV clusters and the RP of inhibitory synapses fits with previous reports that synapsin 1 clusters liposomes *in vitro* (Milovanovic et al., 2018) and that a synapsin/intersectin mixture coacervates with purified SVs (Wu et al., 2020).

In presynaptic terminals, ATP is both synthesized and consumed in an activity-dependent manner (Rangaraju et al., 2014). ATP is known to influence many processes within presynaptic terminals (Brautigam et al., 2004; Shulman et al., 2015). Our work adds synapsin LLPS to this list: we found that ATP inhibits LLPS (Figures 2A and 2B) within the physiological range of ATP concentration (Rangaraju et al., 2014). This could enable the contribution of ATP-sensitive LLPS to trafficking of GABAergic SVs to vary dynamically according to the physiological status of a presynaptic terminal.

### Tetramerization of synapsin and clustering of glutamatergic SVs

Although it has long been known that synapsins oligomerize (Hosaka and Sudhof, 1999; Brautigam et al., 2004; Orlando et al., 2014), the physiological function of synapsin oligomerization was unknown. We have discovered that SVs are densely interconnected under *in vitro* conditions that promote synapsin 2a tetramerization (Figures 2H-2L), an effect that is reduced by synapsin 2a mutants that do not tetramerize (Figures 3D-3G). At excitatory synapses, these mutants do not rescue defects in total SV pool size or RP size (Figures 5C-5E) or mobilization of SVs during synaptic depression (Figures 5H and 5K). Further, perturbing LLPS does not prevent rescue of glutamate release by synapsin 2a (Figures 6G-6J). These results establish a role for synapsin tetramerization in maintaining the RP of glutamatergic SVs. Filamentous structures have long been postulated to connect and cluster SVs (Hirokawa et al., 1989; Siksou et al., 2007; Fernandez-Busnadiego et al., 2010; Cole et al., 2016; Wesseling et al., 2019) and are at least partially dependent on synapsins (Wesseling et al., 2019). Our results have identified synapsin tetramers as potential molecular mediators of these inter-vesicular links.

At excitatory synapses, tetramerization-defective synapsin 2a mutants did not rescue mobilization of SVs from the RP, indicating that tetramerization of synapsins is required both for maintaining glutamatergic SVs in the RP and for mobilizing these SVs. Further work will be needed to determine whether the signals that control glutamatergic SV mobilization – such as Ca^2+^ (Schiebler et al., 1986; Wheeler et al., 1994; Schneggenburger and Neher, 2000) and protein kinases (Greengard et al., 1993; Hosaka et al., 1999; Chi et al., 2003) – work by disassembling synapsin tetramers.

### Two synapsin-dependent clustering mechanisms

Previous work established that synapsins have different functions at GABAergic and glutamatergic synapses (Gitler et al., 2004a; Gitler et al., 2008; Song and Augustine, 2016). Here we have clarified the nature of these functions. At GABAergic synapses, the total pool of GABAergic SVs (Figure 4E) and the kinetics of GABA release from TKO neurons (Figure 4H and 4I) are maintained by synapsin 2a mutants with impaired tetramerization. Although synapsin dimerization is intact in these mutants, rescue is due to LLPS, rather than dimerization: the intersectin SH3A domain maintained dimerization (Figure S6) yet prevented both LLPS and rescue (Figures 6D and 6E). Because every major synapsin isoform has a substantial IDR and is capable of producing LLPS (Figure 1), this can account for the previously paradoxical observation that each isoform can rescue GABA release from synapsin TKO neurons (Song and Augustine, 2016) even though only synapsin 2a rescues glutamate release (Gitler et al., 2008).

Our comparison of RP size and SV mobilization from the RP between GABAergic and glutamatergic synapses established a possible rationale for employing the same molecule, synapsin, to serve two different functions at these two types of synapses (Figures 7A-7F and S7). GABAergic synapses had a smaller RP (Figure S7C) than that of glutamatergic synapses, due to a weaker ability of synapsin LLPS to clustering SVs (Figure 2K). This slows the rate of SV mobilization from the RP to the RRP and additionally lowers the probability of GABA release from the RRP (Figures 7C and 7F). This combination serves as a “brake” that allows synapsin LLPS to sustain transmission at GABAergic synapses during repetitive activity, despite a smaller RP. This mechanism provides a coherent rationale for employing LLPS, rather than tetramerization-dependent SV crosslinking, at GABAergic synapses. It might also explain the poorly-understood, activity-independent short-term depression observed at hippocampal GABAergic synapses (Hefft et al., 2002). On the other hand, the rationale for employing tetramerization-dependent SV crosslinking at glutamatergic synapses is clear: this mechanism provides a greater ability to cluster SVs (Figures 2K and 3E), yielding a larger RP of glutamatergic SVs (Figure S7C) and a faster rate of mobilization of these SVs during repetitive synaptic activity (Figure 7C).

In conclusion, our results provide a striking demonstration of the molecular distinctions between SV trafficking mechanisms at excitatory and inhibitory synapses (Takikawa and Nishimune, 2022) Given the high molecular diversity of synapses (O’Rourke et al., 2012), it remains to be determined whether synapsin tetramerization and LLPS play similar roles at other excitatory and inhibitory synapses, respectively.

## Acknowledgments

We thank K. Chung, P. Teo and M. Yeow for excellent technical and administrative support and T.H. Ch’ng, D. Gitler, F. Meunier and Y. Saheki for helpful comments on our manuscript.

## Author contributions

S.-H.S. and G.J.A. designed research; S.-H.S. performed research; S.-H.S. contributed reagents/analytic tools; S.-H.S. analyzed data; S.-H.S. and G.J.A. wrote the paper.

## Data and materials availability

All data are available in the main text or the supplementary materials.

## EXPERIMENTAL PROCEDURES

### Constructs and molecular cloning

cDNA of rat synapsin isoforms (synapsin 1a: Accession: M27812.1 GI: 206920, synapsin 1b: Accession: M27924.1 GI: 206932, synapsin 2a: Accession: M27925.1 GI: 206833, synapsin 2b: Accession: M27926.1 GI: 206835, synapsin 3a: Accession: AF056704.1 GI: 3170560) were cloned pEGFP-C described in Gitler et al. (*1*). 6XHIS sequence was encoded into N-terminus of pEGFP-C synapsin isoform constructs for protein purification. Synapsin 2a K270Q mutant and 2a K337Q mutant were generated using site-directed mutagenesis method (for K270Q, forward: 5’-GTTTCCTGTCGTGGTGCAGATTGGCCATGCTCAC-3’, reverse: 5’-GTGAGCATGGCCAATCTGCACCACGACAGGAAAC-3’; For K337Q, forward: 5’-AGGGAACTGGCAGACAAACACTG-3’, reverse: 5’-GAGATGGATGTCCTCATG-3’).

### Microisland neuron culture

Homozygous synapsin TKO mice were derived by serial breeding of the mice described in Gitler et al. (2004a). Microisland cultures of hippocampal neurons were prepared as described previously (Nishiki and Augustine, 2004), Glial microislands were prepared to support neuronal survival before seeding dissociated hippocampal neurons. Hippocampus were obtained from TKO newborn pups (postnatal days 0 –2) and dissociated hippocampal neurons were seeded onto glial microisland (Gitler et al., 2004a). Neurons were allowed to mature for 12–18 d before being used for electrophysiological recording.

### Lentiviral vectors

Synapsins were subcloned from pEGFP vectors (Gitler et al., 2004b) into the pFUGW plasmid (Lois et al., 2002), which was constructed by inserting the following into the multicloning site of HR’CS-G (vector backbone from I. Verma, Salk Institute): HIV-1 flap sequence, amplified by PCR from the HIV NLA4.3 genome, the human polyubiquitin promoter-C (gift from L. Thiel, Amgen), the EGFP gene, and the Woodchuck hepatitis virus posttranscriptional regulatory element WRE. The inserted genes were confirmed via sequencing. Lentivirus was prepared as described in ref. *51*. Neurons were infected 2–4 d after plating and analyzed 10-14 d after infection and expression of synapsin isoforms. These neurons were then examined as in ref. *34*.

### Co-immunoprecipitation and Western blotting

GFP- or Flag-synapsin constructs were separately transfected into HEK 293 cells using Lipofectamine 2000 (Promega). Transiently transfected cells were washed with PBS and lysed in immunoprecipitation buffer (20 mM HEPES, pH7.4, 0.5% Triton X100, 150 mM NaCl, 1mM EDTA, 1 mM EGTA, 100 mM NaF, 10 mM Sodium pyrophosphate, 1 mM sodium vanadate, 1 μg/ml leupeptin, 1 μg/ml aprotinin and 1 μg/ml pepstatin). Lysates were clarified by centrifugation at 22,500 g for 10 min. Appropriate antibodies were incubated with clarified lysate for overnight at 4°C and protein G sepharose beads (GE Healthcare) were incubated for 2h at 4°C. The immunoprecipitates were washed three times with immunoprecipitation buffer to remove unbound proteins. Proteins bound to antibodies were loaded in Laemmli sample buffer, separated by SDS-PAGE and immunoblotted. The membrane were incubated in primary antibody for overnight at 4 °C and secondary antibody was incubated for 1 h at room temperature. Antibodies used: rabbit anti-synapsin 2a (G281) was gifted from the Paul Greenguard Lab. mouse anti-synaptiphysin (sc-17750, Santa Cruz), mouse anti-PSD 95 (sc-32290, Santa Cruz), mouse anti-synaptobrevin 2 (104 211, Synaptic Systems), mouse anti-sodium potassium ATPase (ab7671, Abcam) rabbit anti-succinate dehydrogenase (MA5-32598, Thermo Fisher).

### Purification of recombinant synapsin proteins

Expi293 cells (Thermo Fisher Scientifics) were used for expression of 6Xhis-tag constructs according to manufacturer’s manual. After 3-4day expression of proteins, cells were harvested and lysed in M-per mammalian protein extraction reagent (Thermo scientific) with protease inhibitor (EDTA-free, Nacalai) and 10 mM β-mercaptoethanol. The lysates were clarified 14,000 g for 15 min. the clarified lysates were incubated with Ni-NTA magnetic agarose beads for 60 mins in equilibration buffer (50 mM sodium phosphate, 0.3 M NaCl, 10 mM imidazole, 0.05% Tween-20, pH8.0) and Ni-NTA magnetic agarose beads were washed three times with wash buffer (50 mM sodium phosphate, 0.3 M NaCl, 15 mM imidazole, 0.05% Tween-20, pH8.0). His-tag proteins were eluted with elution buffer (50 mM sodium phosphate, 0.3 M NaCl, 0.3 M mM imidazole, pH8.0). Purified proteins were dialyzed overnight with dialysis buffer (25mM Tris-HCl, 150 mM NaCl) and stored in presence of 0.5 mM TCEP.

### Measurement of Synapsin 2a oligomerization

His tagged synapsin 2a was purified and dialyzed in 200 mM NaCl, 20 mM NaPO4, 0.1 mM EGTA, pH 7.4. After quantification, 300 nM of synapsin 2a was incubated with 0.5 mM DSS (disuccinimidyl suberate) dissolved in DMSO or with DMSO alone in the with or without ATP (5 mM) and/or Ca^2+^ (2.1 mM) for 1 h at room temperature. The reaction was stopped by the addition of 10 mM Tris, pH 7.4, and the cross-linked synapsin 2a complexes were resolved by SDS-PAGE on 6% (w/v) polyacrylamide gels, transferred to PDVF membranes, and detected by immunoblotting with polyclonal anti-Syn 2a antibodies (G281, generously presented by Paul Greengard’s lab).

### Purification of synaptic vesicles

Synapsin-free synaptic vesicles were prepared from brains of synapsin TKO mice as described in ref. (Huttner et al., 1983). Forebrains were homogenized in Buffered Sucrose (BS; 320 mM sucrose, 4 mM HEPES (pH 7.4), 1 mM EDTA, and 0.25 m M DTT) and the synaptosomal fraction was obtained by a serial centrifugation and hypotonic lysis of the homogenate. As a final step, synaptic vesicles were purified by discontinued sucrose-gradient centrifugation at the 200 to 400 mM interface. Purified synaptic vesicle fraction were dialyzed with the buffer for the next experiment for overnight at 4°C. Crude synaptosomal fraction and purified synaptic vesicles from TWT or TKO were tested by Western blotting (Extended Data Fig. 2b).

### Reconstituting synaptic vesicle clustering

Synaptic vesicles (purified from TKO mice) were labeled with Dil dye (1,1’-dioctadecyl- 3,3,3’,3’-tetramethylindocarbocyanine, Invitrogen; 1 µM) for 1 h at 22°C and dialyzed with the buffer containing 150 mM NaCl, 25mM Tris pH 7.4 for overnight at 4°C. Purified EGFP-synapsin 2a was also dialyzed in the same buffer, with TCEP (0.5 mM final concentration) added.

75 ng/μl of synaptic vesicle and 10 uM of EGFP synapsin 2a was incubated in the buffer contained 25 mM Tris-HCl (pH 7.4), 150 mM NaCl and 0.5 mM TCEP with 3% PEG. The final mixture was placed in cover glass which is coated poly-D-lysine and observed under the confocal microscope. Images were collected with a 60X 1.2NA water-immersion objective (Olympus).

### Synaptic vesicle binding

To measure binding of synapsins to purified synaptic vesicles, 5 ug of synaptic vesicles were mixed with purified synapsin protein in SV binding buffer conditioned 0.25 mM glycine, 30 mM sucrose, 30 mM NaCl, 5 mM Tris-Cl, 4 mM HEPES (pH 7.4) 0.02 EDTA, 0.22 NaN3, and 100 µg/ml BSA. and incubated for 1 hour on ice. The mixture was floated 100 ul of 10% sucrose and ultra-centrifuged for 30 minutes at 220,000Xg. To detect co-sedimented synapsin proteins, the sedimented synaptic vesicles were analyzed by immunoblotting with an anti-synapsin antibody.

### FM dye staining and post-hoc immunostaining

Cultured neurons were incubated for 1 min with fluorescent dye N-(3-triethylammonium-propyl)-4-(6-(4-diethylamino)phenyl)-hexatrienyl)pyridinium dibromide (FM4-64; 4 µM; Invitrogen) and loaded by stimulation (10 Hz, 1200 AP). The saline used for FM dye contains CNQX (10 µM) and AP5 (50 µM). After additional 2 min incubation in FM4-64 containing saline, neurons were washed with FM dye free saline for 40 min and imaged with stimulation as described in Results. To distinguish excitatory and inhibitory synapse, neurons were stained with vesicular glutamate transporter antibody (VGLUT, Thermo Fisher) or with vesicular GABA transporter (VGAT, Thermo Fisher). Boutons which were coloclized with VGLUT antibody (135 303, synaptic system) or VGAT antibody (131 003, Synaptic Systems) as well as expressing GFP-synapsin, were selected and their fluorescence signal was analyzed using ImageJ (NIH). The background was subtracted to reduce the deviation across the population of bouton. Cultured neurons for FM day staining were used from at least three different culture set.

### Imaging and image analysis

Protein without or with synaptic vesicle were mixed and incubated in 25 mM Tris-HCl (pH 7.4), 150 mM NaCl, 0.5 mM TCEP. The mixture was placed on 35 mm glass bottom confocal dish (SPL life science) and covered with cover slip. Glass bottom was coated with poly-D-lysine (Sigma-Aldrich) before use. Imaging was performed using a laser-scanning microscope (FV1000MPE; Olympus) equipped with ×60 NA 1.1 (Olympus LUMFLN) water-immersion objectives lens. Excitation wavelengths were 488 nm for EGFP and 591 nm for Dil. All images were analyzed with ImageJ (NIH). Images of EGFP-synapsin droplets were acquired by z-stack of 3 slices (0.3 µm interval) and the background was subtracted by the same value of rolling ball radius method. Droplets were detected by thresholding the image. The threshold was determined by the same auto-threshold algorism for each condition. Size and intensity were measured by the ‘analyze particle’ function of ImageJ. Synaptic vesicle clustering was also measured in the same way.

### Electrophysiology

Whole-cell patch-clamp recordings were made from cultured hippocampal neurons, as described in ref. *31*. Patch pipettes (4-7 MΩ) were filled with intracellular solution containing (in mM): 50 K-glutamate, 71 K-gluconate, 15 NaCl, 6 MgCl_2_, 2 EGTA, 2 Na_2_ATP, 0.3 Na_2_GTP, 20 HEPES-KOH (pH 7.4, adjusted with KOH, 285 mOsm). The extracellular solution contained (in mM): 150 NaCl, 3 KCl, 2 CaCl_2_, 2 MgCl_2_, 20 D-glucose and 10 HEPES-NaOH, pH 7.3 (310 mOsm). Neurons were voltage clamped at -70 mV with a Multiclamp 700B amplifier (Molecular Devices) and a Digidata 1440 interface (Molecular Devices). All recordings were performed at room temperature (21-25°C). Electrophysiological data were sampled at 10 kHz and filtered at 10 kHz. Axonal action potentials were evoked by depolarizing the cell body to +40 mV for 0.5 msec every 15 sec. To minimize series resistance errors (Marty and Neher, 1995), only recordings with a series resistance <20 MΩ were analyzed. To account for potential effects of these constructs on EPSC kinetics(Hilfiker et al., 1998; Humeau et al., 2001; Hilfiker et al., 2005; Medrihan et al., 2013; Song and Augustine, 2016). we measured the charge of each EPSC, converted it to quantal content, and then integrated quantal content over the entire stimulus train (Fig. 6D).

### Statistical analysis

Data are expressed as means ± SEM. All statical analysis was performed using Origin (OriginLab) or Prism 7 (GraphPad) software. Normality test for data by Kolmogorov-Smirnov test confirmed that all data were normally distributed. Normally distributed data was compared using the Students t-test, or one-way ANOVA with Tukey’s post hoc analysis in the case of multiple comparison.

## Supplementary figure legends

**Figure S1.**
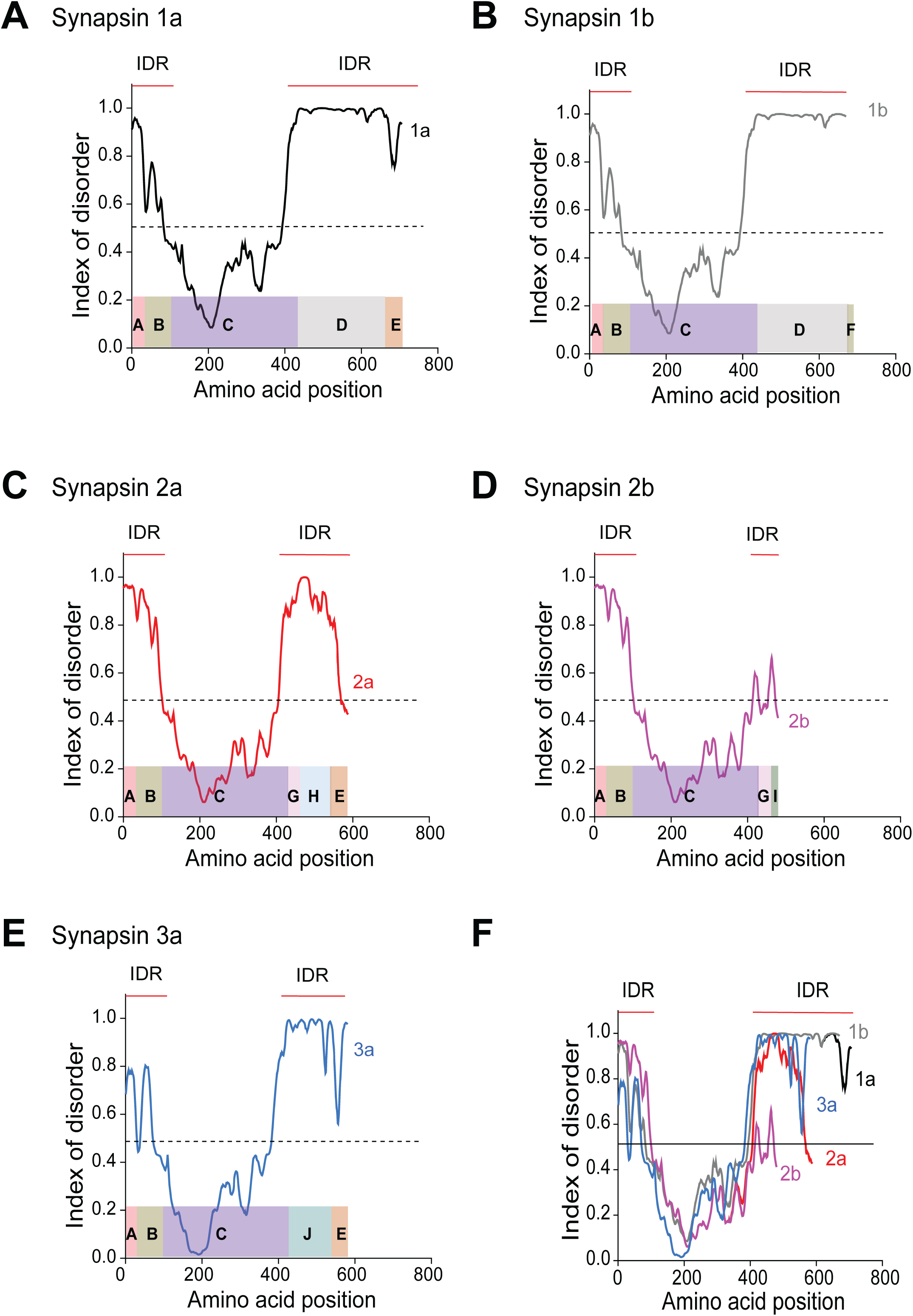
Comparison of intrinsically disordered region of synapsin isoforms. (A-E) Structure of intrinsically disordered regions (IDRs) of indicated synapsin isoforms. (F) Comparison of IDRs of all 5 synapsin isoforms.

**Figure S2.**
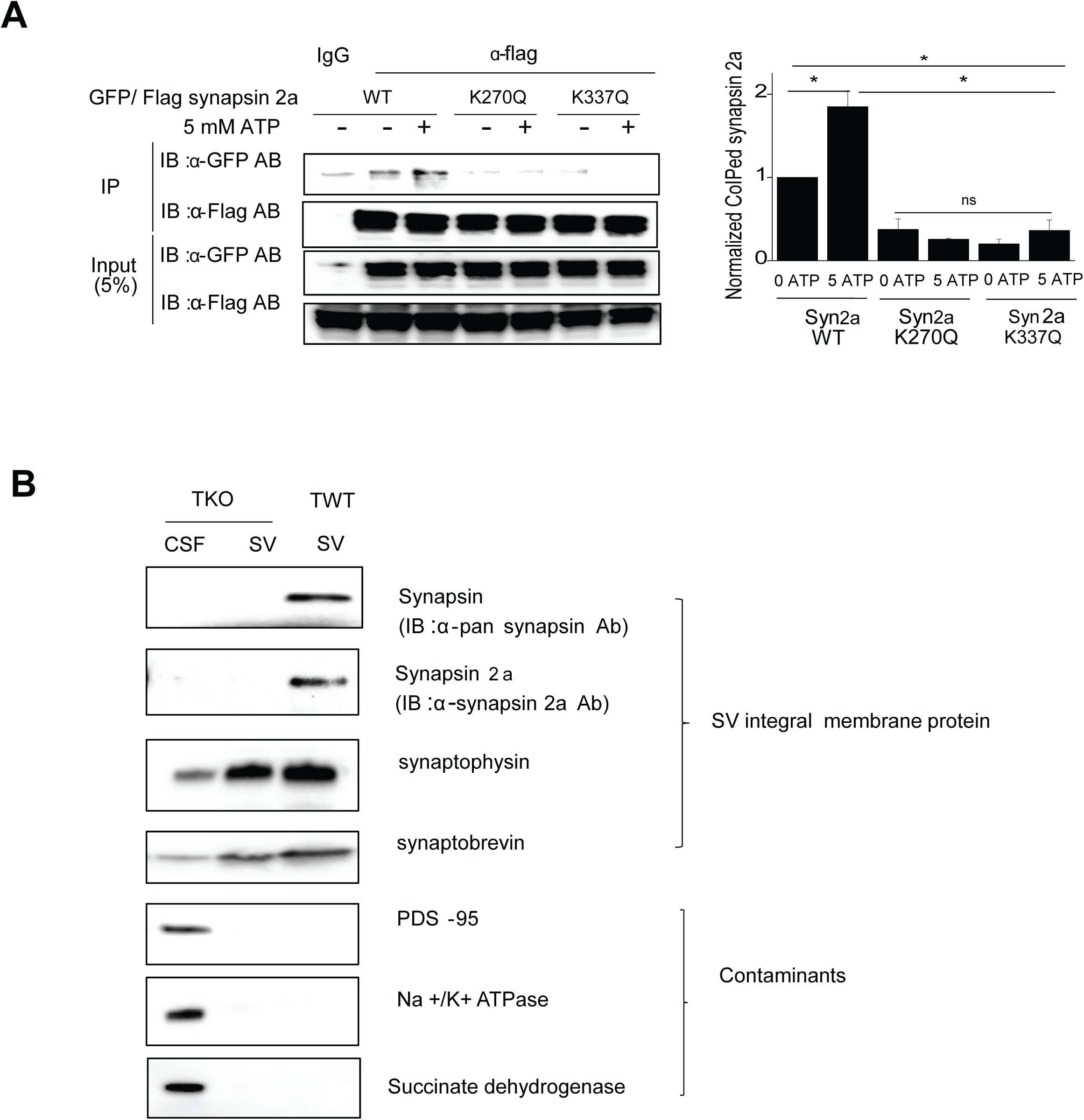
Oligomerization of synapsin 2a and SV purification. (A) Oligomerization defects in synapsin 2a mutant K270Q and K337Q proteins expressed in HEK 293T cells. Western blots (*left*) and quantification (*right*) indicate reduction in both ATP-dependent and ATP-independent association. Comparisons were done with ANOVA, followed by Tukey’s post-hoc test; asterisks indicate significant differences (p<0.05; n=4). (B) Immunoblot characterization of purity of crude synaptosomal fraction (CSF) and purified synaptic vesicle fraction (SV) from TKO and TWT mouse brains.

**Figure S3.**
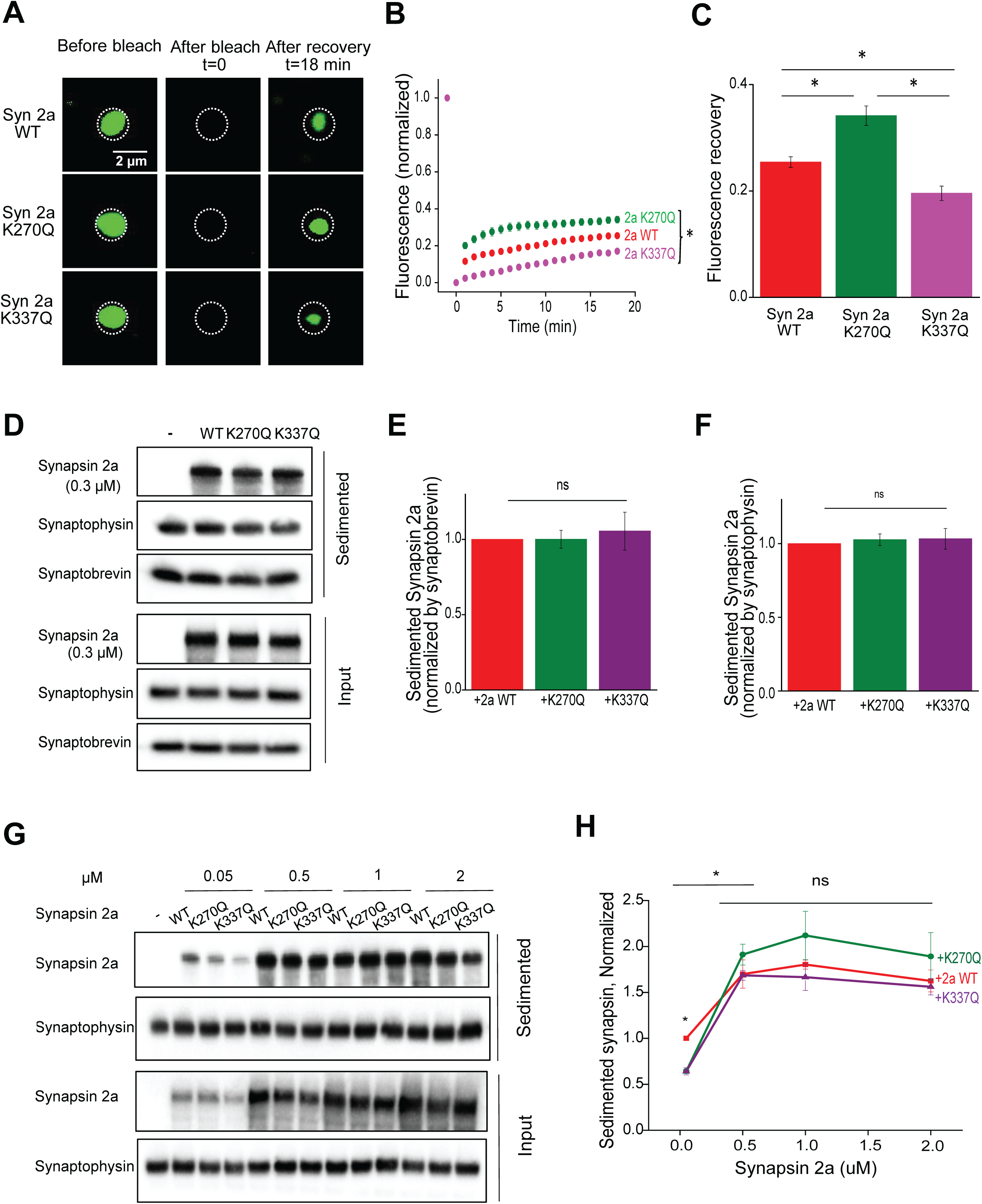
FRAP of synapsin 2a and SV binding. (A) Images of LLPS-dependent droplets before and after photobleaching for 3 synapsin 2a variants. (B) Time course of fluorescence recovery after bleaching for each synapsin 2a variant. (C) Mean rates of fluorescence recovery, measured 20 min after photobleaching, from experiments shown in (B), Statistical comparisons in (C) were done with ANOVA, followed by Tukey’s post-hoc test; asterisks indicate significant differences (p<0.05). Number of replicates ranged from 13-17. (D-F) Binding of synapsin 2a variants to SVs. Purified SVs from TKO mice were mixed with purified synapsin 2a protein variants and sedimented. (D) Immunoblotting indicated that all 3 synapsin 2a variants co-sediment with SVs. (E-F) Co-sedimentation of synapsin variants with SVs, measured either by synaptobrevin-2 (E) or synaptophysin (F). Statistical comparisons were done with ANOVA, followed by Tukey’s post-hoc test; asterisks indicate significant differences (p<0.05; n=4). (G-H) Concentration-dependent binding of purified synapsin 2a protein variants to purified SVs. (G) Immunoblot analysis of co-sedimentation of synapsins with SVs. (H) Quantification of data in (G); all 3 synapsin variants bound to SVs with similar concentration dependence. Statistical comparisons were done with ANOVA, followed by Tukey’s post-hoc test; asterisks indicate significant differences (p<0.05; n=5).

**Figure S4.**
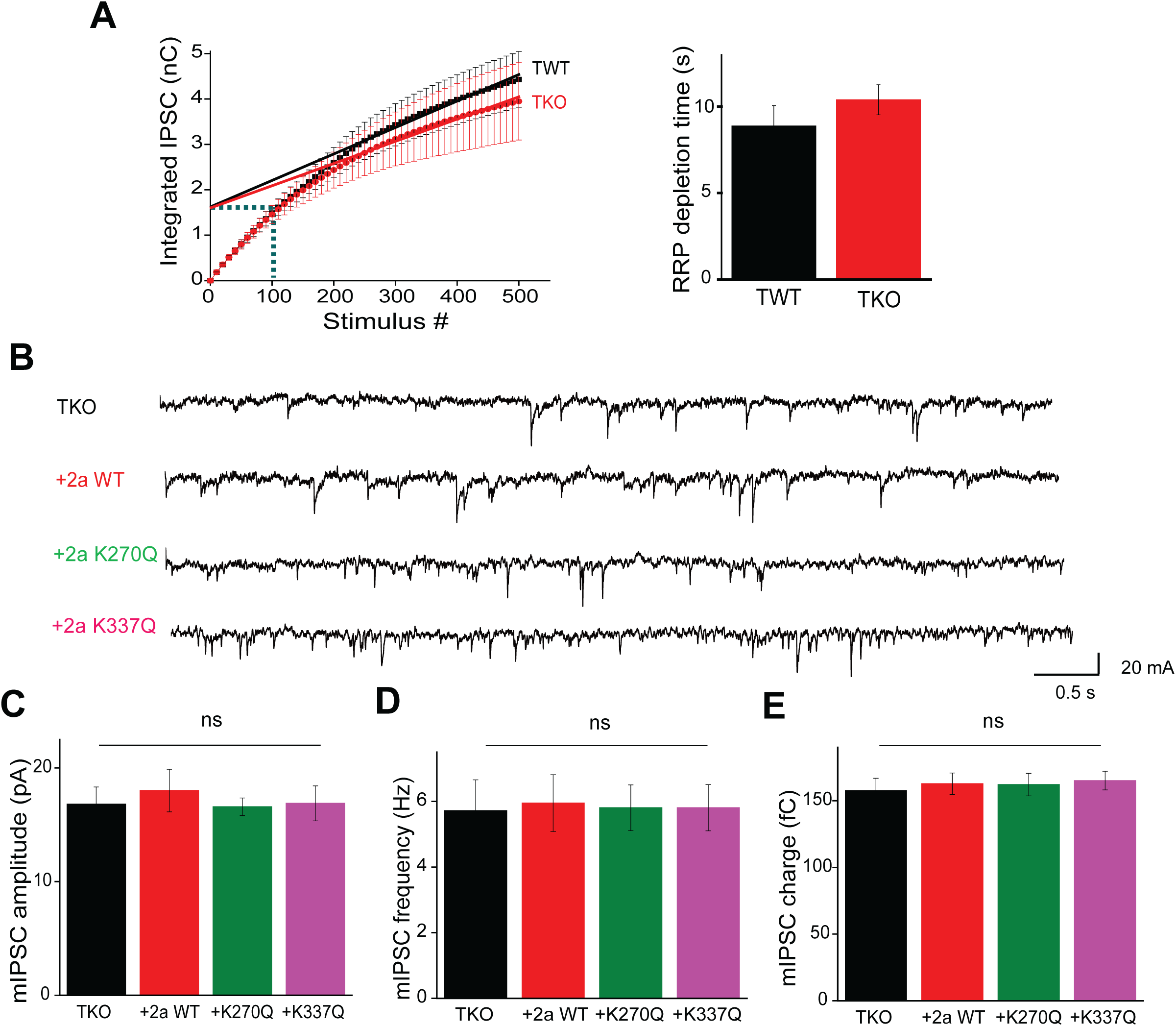
Time course of RRP depletion in GABAergic synapses and mini IPSCs from synapsin 2a and mutants. (A) *Left,* Time course of cumulative IPSC charge during trains of 500 stimuli (10 Hz). RRP size was estimated by extrapolation by linear regression fits to data from 20-50 s. Dotted lines indicate time required for RRP depletion. *Right*, mean values of time required for RRP depletion in TKO and wild-type (TWT) neurons. (B) Representative traces of spontaneous miniature IPSCs (mIPSCs) recorded in TKO neurons and TKO neurons expressing each synapsin variant. (C-E) Analysis of mIPSC properties: mean amplitude (C), mean frequency (D) and mean mIPSC charge (E). Statistical comparisons were done with ANOVA, followed by Tukey’s post-hoc test; no significant (p<0.05) differences were observed. Number of experiments ranged from 14 to 17.

**Figure S5.**
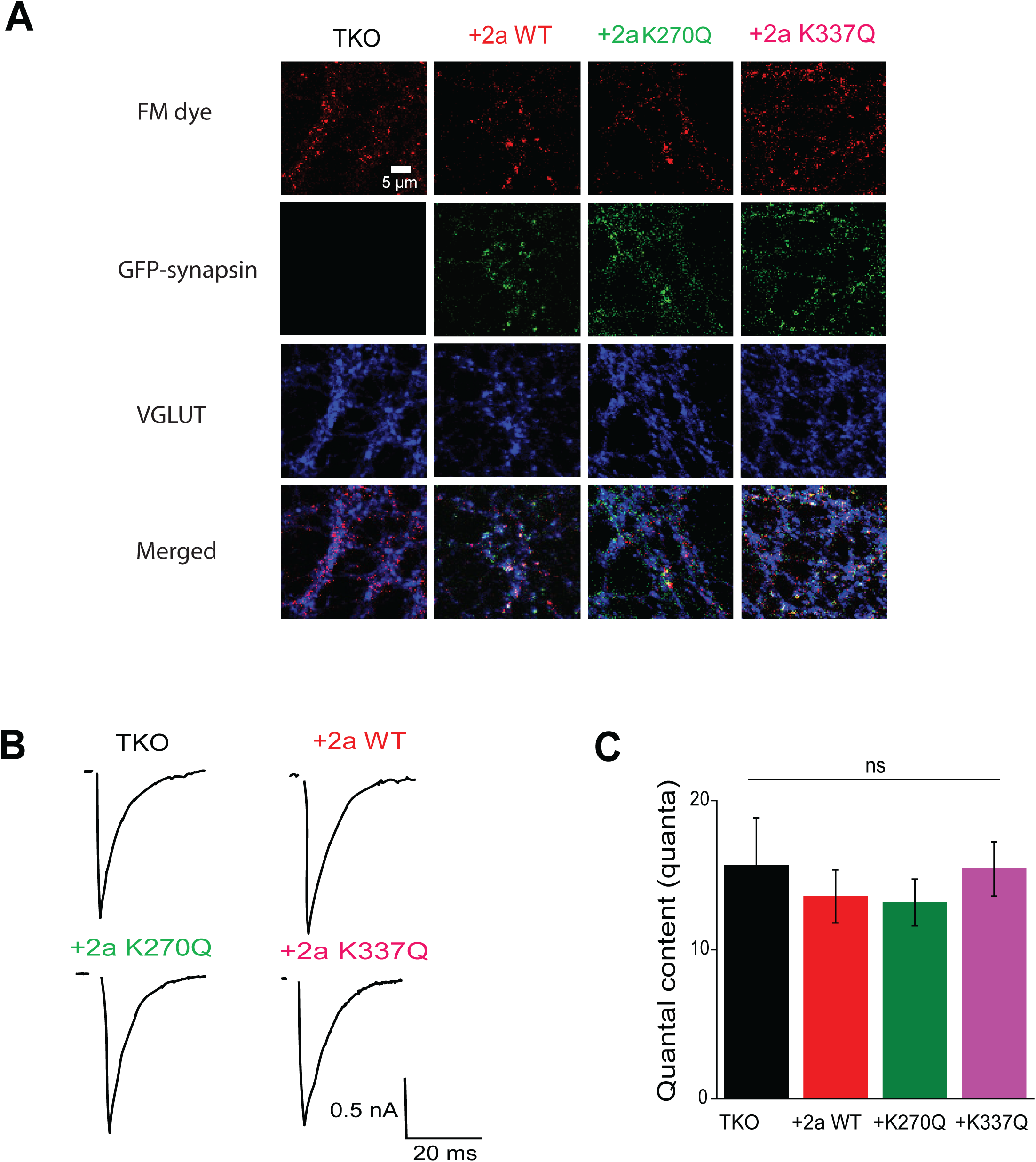
Effects of tetramerization at glutamatergic synapses. (A) Representative images of FM4-64 dye loading (top row) of presynaptic terminals expressing GFP-synapsin 2a (second row). Post-hoc immunostaining for VGLUT (third row) indicates glutamatergic presynaptic terminals. Bottom row is merger of images in 3 rows above, to compare co-localization of fluorescent signals. (B) Representative EPSCs evoked by single presynaptic action potentials in TKO neurons not expressing any synapsins or expressing the indicated synapsin 2a variants. (C) Mean EPSC quantal content in the indicated conditions. Statistical comparisons were done with ANOVA, followed by Tukey’s post-hoc test; no significant (p<0.05) differences were observed. Number of experiments ranged from 14 to 17.

**Figure S6.**
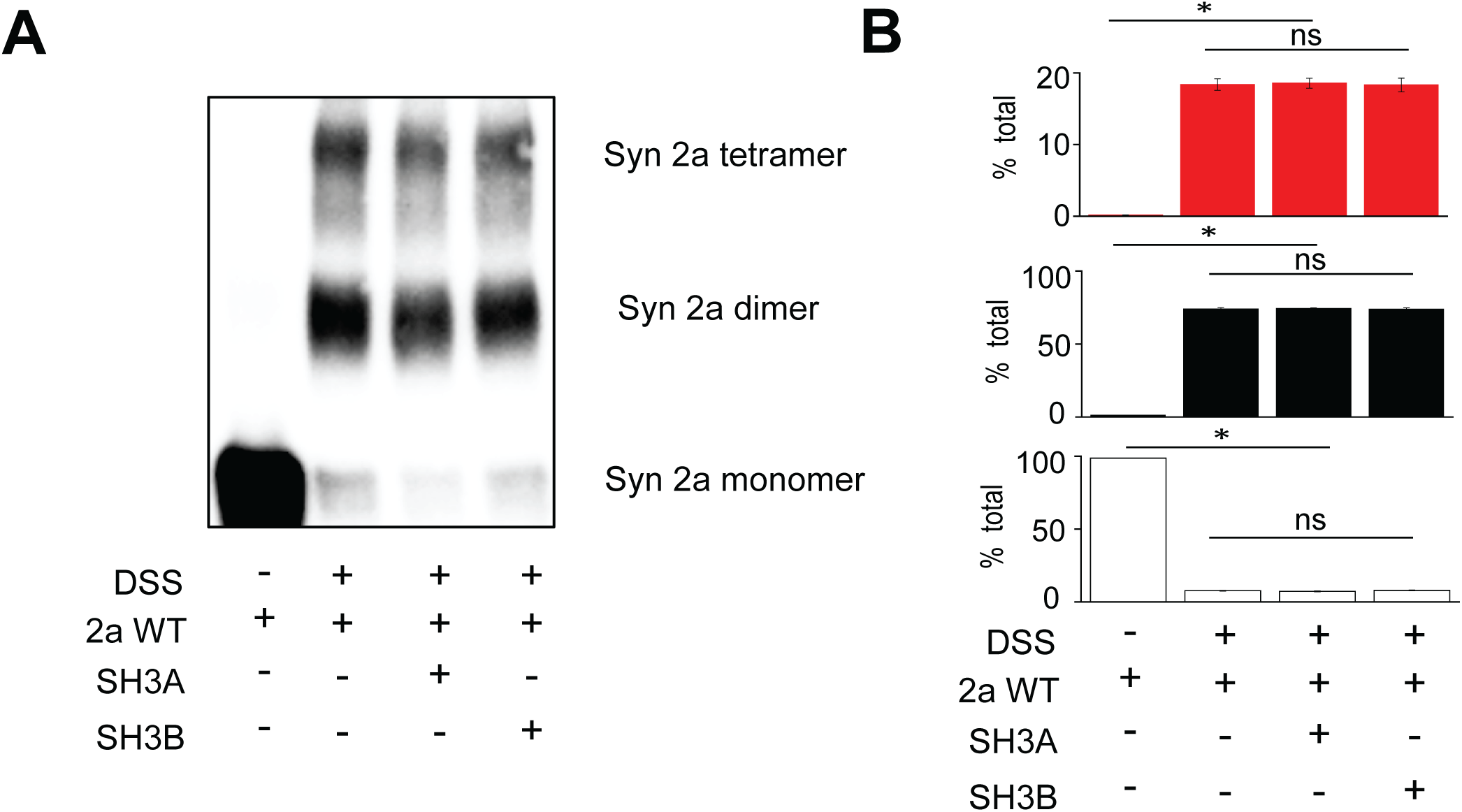
Effect of SH3 domains on synapsin 2a oligomerization. (A) Western blot analysis of synapsin 2a oligomerization with and without intersectin SH3A or SH3B domains, in the presence of 5 mM ATP and crosslinking reagent, DSS. (B) Quantification of results in (A) indicate no effect of SH3 domains on synapsin 2a. Statistical comparisons were done with ANOVA, followed by Tukey’s post-hoc test; no significant (p<0.05) differences were observed. Number of independent replicates is 4.

**Figure S7.**
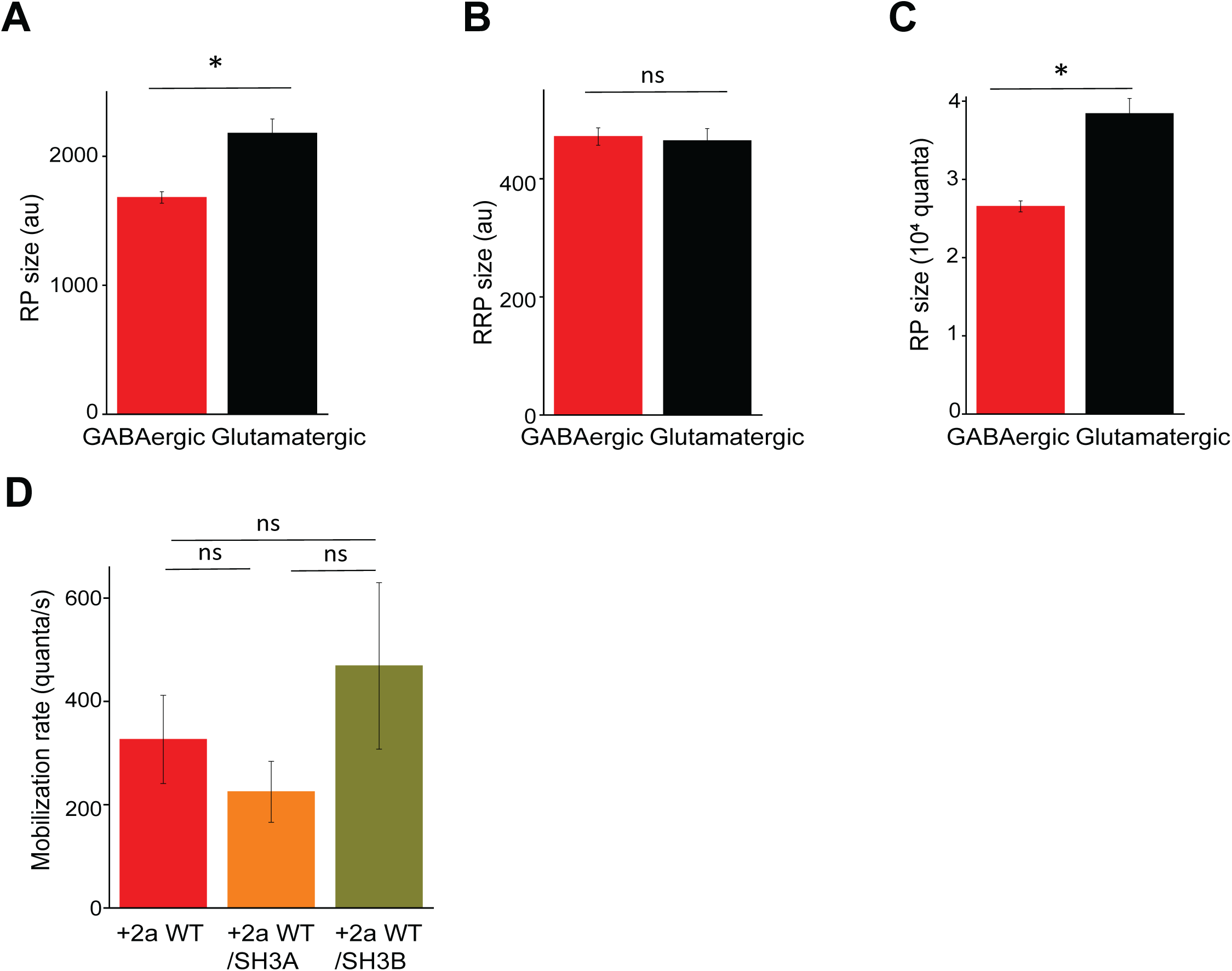
Determination of number of quanta in the RP of GABAergic and glutamatergic synapses. Comparison of RP size (A) and RRP (B) size between GABAergic and glutamatergic synapses, based on FM4-64 dye experiments. (C) Comparison of number of quanta in the RP. The amount of FM dye labelling per quantum was measured by dividing RRP size measured in FM4-64 dye experiments (B) by the number of quanta in the RRP, determined from cumulative number of quanta released during trains of stimuli, as measured electrophysiologically (Figure 7B). (D) Comparison of mean mobilization rates during train stimulation (Figure 7D), which did not show any significant differences between groups. For (A-C), statistical comparisons were done with a t-test; significant (p<0.05) differences are indicated by asterisks, while *ns* indicates non-significant differences. were observed in (A) and (C). For (D), statistical comparison was done by ANOVA with tukey’s post hoc test. Number of experiments are indicated in legends of Figures 4, 5 and 7.

